# Full-length next-generation sequencing of 11 HLA loci of more than 1000 individuals from clinical cohorts in East and West Africa

**DOI:** 10.1101/2025.09.04.673795

**Authors:** Yvonne V. Rosario, Aviva Geretz, Lakshmi Rani Iyer, Philip K. Ehrenberg, Alýa Tyson, Hannah Kibuuka, Fred Wabwire-Mangen, Lucas Maganga, Abdulwasiu Tiamiyu, Jonah Maswai, Josphat Kosgei, Frederick Sawe, Gary R. Matyas, Merlin L. Robb, Julie A. Ake, Rasmi Thomas

## Abstract

Human Leukocyte Antigen (HLA) loci have been implicated in several diseases from different world populations, including HIV-1. It is necessary to characterize HLA allele variation at the population level prior to investigating associations linked to human diseases. In global databases, limited high-resolution HLA allele types generated by next-generation sequencing (NGS) have been described for populations from African countries. We sought to expand our HLA NGS database to include a total of 1023 participants from multiple HIV clinical studies using full-length HLA genotyping by NGS. Collectively we describe HLA genotypes of individuals from Kenya (n=375), Uganda (n=338), Nigeria (n=139), Tanzania (n=89), and Mozambique (n=82). Overall, we identified 371 unique HLA alleles across 11 loci with the most frequent at each locus being A*02:01:01, B*53:01:01, C*04:01:01, C*06:02:01, DPA1*01:03:01, DPB1*01:01:01, DQA1*01:02:01, DQB1*06:02:01, DRB1*15:03:01, DRB3*02:02:01, DRB4*01:03:01, and DRB5*01:01:01. A total of 25 novel alleles were identified, including 4 with non-synonymous changes affecting the peptide binding groove of HLA molecules. This expansion of NGS based HLA data at the African population level will improve our understanding of human genetic variation and provide insights for vaccine development and targeted personalized therapies.

## 1. Introduction

Human Leukocyte Antigen (HLA) are highly polymorphic immunoreceptor proteins derived from chromosome 6p21 within the Major Histocompatibility Complex (MHC) region of the human genome^1,2^. HLA plays a key role in peptide antigen presentation during host immune responses to microorganisms including human immunodeficiency virus type 1 (HIV-1), and several studies have defined locus-specific associations^3-8^. The two major classical HLA are class I (HLA-A, B, C) and class II (HLA-DP, DQ and DR)^1^. During HIV-1 infection, HLA class I immunoreceptors on infected nucleated cells present HIV-1 derived peptides, enabling cytotoxic CD8+ T cell surveillance, recognition, and elimination of infected cells^9,10^. Heterozygosity within HLA class I offers greater functional divergence between allotype pairs, thus improving epitope binding site fitness, and is associated with slower disease progression^11,12^. HLA class II molecules present exogenous HIV-1 peptides to CD4+ T cells for activation and canonical immune responses that help B cells make antibodies.

Although the HIV epidemic is in decline, more than 25.6 million people living with HIV reside in Africa^13^. As HLA surface proteins enable the critical function of antigen presentation, greater allelic variation offers greater population-wide and individual fitness against various pathogens^11^. HIV-1, the most common type of HIV, has diversified into multiple subtypes, which exhibit different levels of adaptation to the human immune system^14^. Understanding HIV adaptation is crucial for developing effective vaccines and therapies, as vaccines ideally need to target conserved viral epitopes to minimize potential for immune escape. The inherent infidelity of the viral reverse transcriptase, together with the absence of adequate repair mechanisms, favors generation of viral variants that adapt to environmental conditions, particularly virus-specific immune responses^15^.

Different ethnic groups have varying HLA allele frequencies, and HIV strains are evolving to evade immune responses specific to those HLA profiles^16,17^. In particular, African populations exhibit the greatest diversity of HLA alleles globally, compared to other ethnic groups^18-20^, concomitant with many circulating HIV-1 subtypes, with subtype A being prevalent in East Africa, while CRFO2_AG is dominant in West Africa^21^. Virus adaptation can influence how quickly HIV progresses to AIDS, with some HLA alleles being associated with disease outcomes. Population-specific HLA class I associations with HIV-1 disease protection or progression were first identified in the mid-1990s and have continued to be characterized over the course of the HIV-1 epidemic, showing that novel HLA associations likely remain to be discovered^6,22-29^. Understanding HLA class I restricted immune responses is crucial because HLA drives viral evolution through immune escape, impacting disease pathogenesis^30^. Similarly, HLA class II molecules present peptides to CD4+ T cells, which can then induce B cells to produce antibodies. Interestingly, the influence of specific class II alleles on antibody production has been shown to affect HIV-1 vaccine efficacy^30^. Therefore systematic characterization of HLA in different populations is necessary because HLA alleles are distinct and vary by geographical location^16,31,32^.

High resolution HLA typing generated by Next-Generation Sequencing (NGS) offers increased accuracy and scalability over low-to-medium resolution approaches, which will improve targeted vaccine development and immunologic studies^33-35^. While advances in NGS and third generation sequencing technologies have driven the rapid expansion of HLA allele discovery, only limited high-resolution HLA allele typing has been performed in individuals from African countries^31^. Exploring additional genotype profiles from African cohorts more deeply is necessary to characterize unique population alleles and uncover novel or rare HLA alleles with potential roles in HIV disease progression and/or protection. We previously developed a population agnostic HLA genotyping method to characterize world populations where clinical trials are conducted^30,33-39^.

This method was used to describe full-length NGS HLA allele and haplotype frequencies from Thai and Philippine populations, which allowed for identification of unique, novel, and rare alleles^34,35^. This enabled identification of alleles that associated with vaccine efficacy and disease progression in Thailand^28,30^. Here we implement this method to characterize HLA from 11 loci (HLA-A, B, C, DPA1, DPB1, DQA1, DQB1, DRB1, DRB3/4/5) from more than 1000 individuals from clinical trials conducted in Kenya, Uganda, Tanzania, Mozambique, and Nigeria. The goal was to define high-resolution HLA genotypes across different countries in Africa, with a sample size greater than 50 from each country, and to assess if allele frequencies were associated with geographic location. Our findings indicate a high degree of allelic diversity within and between the different African countries when compared to other global populations. These findings will be key to understanding immune responses after infection and vaccination in continental African populations.

## 2. Methods and materials

### 2.1 Study Populations

Peripheral blood mononuclear cells (PBMCs) were collected for HLA typing from 1023 individuals from Kenya, Uganda, Tanzania, Mozambique, and Nigeria (Fig. 1A). Participants were enrolled in one of the following vaccine trial cohorts: RV217 (n=59), RV251 (n=187), RV329 (n=165), RV456 (n=488), or RV460 (n=124) (Fig. 1C). RV217 was a multisite study in East Africa and Thailand that enrolled HIV negative individuals at high risk to observe and track sero-status^40^. Participants with changes in HIV status were characterized during acute HIV infection. RV251 (NCT01549470) was a Phase 1 HIV vaccine clinical trial conducted in Kampala, Uganda. Adults negative for HIV were enrolled and monitored for immunological changes in response to multiclade HIV-1 DNA plasmid vaccine VRC-HIVDNA009-00-VP^41^. The RV329 African Cohort Study (AFRICOS) is a long-term HIV observational cohort study in Kenya, Uganda, Tanzania, and Nigeria ^42^. RV456 (NCT02598388) is a Phase 2 Ebola vaccine multisite study with sites in East and West Africa^43,44^. People without HIV (PWOH) and people living with HIV (PLWH) were enrolled into accelerated 14- or 28-day Ebola vaccine regiments. PLWH were monitored for changes in CD4+ cell counts and viral load. RV460 (NCT04826094) was a Phase 1 HIV vaccine trial conducted in healthy HIV uninfected adults in Kenya. This vaccine study evaluated the safety and efficacy of DNA priming with DNA encoding HIV-1 gp120 envelope protein with or without boosting with an adjuvanted recombinant HIV-1 gp145 protein^45^. HLA genotyping was approved for all sites by the participating local institutional review boards and in the USA.

**Figure 1.**
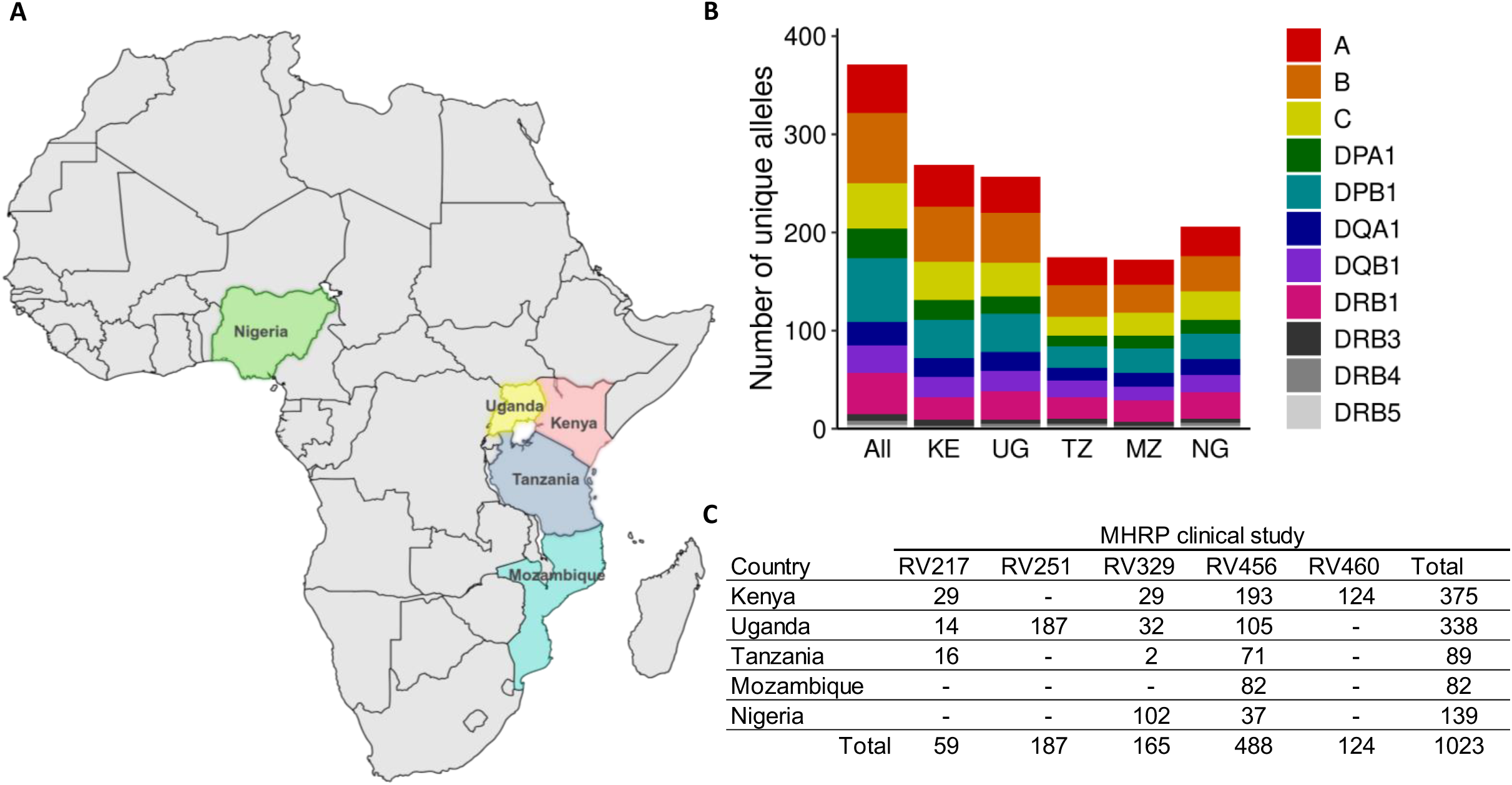
HLA genotyping in 1023 participants from multiple African countries. **A.** Cohort map of Africa displays countries represented in this study across Kenya (shaded pink), Uganda (shaded yellow), Tanzania (shaded blue), Mozambique (shaded teal) and Nigeria (shaded green). **B.** Number of HLA alleles found per locus in all African countries (All), Kenya (KE), Uganda (UG), Tanzania (TZ), Mozambique (MZ), and Nigeria (NG) displayed in stacked bar graphs. **C.** Peripheral Blood Mononuclear Cells (PBMC) were collected from participants enrolled in MHRP clinical studies from RV217 (n=59), RV251 (n=187), RV329 (n=165), RV456 (n=488), and RV460 (n=124).

### 2.2. DNA Extractions

Genomic DNA was extracted from 5 million PBMC using either column-based purifications with the Qiagen DNA Blood Mini kit, or automation with the QIAsymphony and Qiagen DNA Midi kit, per manufacturer’s instructions (Qiagen, Valencia, CA). Quantitation of genomic DNA was completed using a NanoDrop 8000 (Thermo Fisher Scientific, Waltham, MA).

### 2.3 HLA Genotyping

#### 2.3.1 PCR amplification

Contiguous full-length PCR amplifications of up to 11 HLA loci (HLA-A, B, C, DPA1, DPB1, DQA1, DQB1, DRB1 and DRB3/4/5) were performed on 1023 participants using a method developed by our lab^34-37,39^. One microliter from each post-PCR product was screened on a 1% agarose gel, and the remainder was purified with AMPure XP beads using the Biomek NXp Laboratory Automated Workstation (both Beckman Coulter; Brea, CA). The purified amplicons were quantitated with the Quant-iT dsDNA High Sensitivity Assay Kit (Thermo Fisher Scientific) using the Filter Max F3 plate reader with SoftMax Pro 7 software (Molecular Devices; San Jose, CA).

#### 2.3.2. Library Preparation and Sequencing

For each participant, full-length amplicons from up to 11 loci were combined into single 30 µl equimolar pools of 10-20 ng product within designated wells of a twin.tec LoBind PCR plate (Eppendorf; Hamburg, Germany). These pools were subjected to tagmentation using Illumina bead-linked transposome technologies as we describe previously^34-37,39^. Briefly, sequencing libraries were prepared manually or with automation on the epMotion 5075 liquid handler (Eppendorf) using one of the following sequence/index kit combinations: Nextera XT DNA Library Prep with Nextera XT indices; Nextera DNA Flex Library Prep with Nextera DNA CD indices; DNA Prep with Nextera DNA CD indices or DNA/RNA indices sets A-D (all Illumina; San Diego, CA), as per manufacturer’s recommendations. Uniquely dual indexed donor libraries were sequenced on MiSeq or NextSeq 1000 sequencers (both Illumina). For sequencing on the MiSeq, 10 ng inputs of each of the 93 donor libraries along with 2 positive and 1 negative controls were pooled into a single tube. This pooled sample library was then diluted to a 12 pM concentration in HT-1 reagent/5 mM Tris-HCl pH 7.5, spiked with 5% PhiX, and loaded into a MiSeq v2 500 bp (for libraries prepared with Nextera XT) or a MiSeq v3 600 bp paired-end cartridge (for libraries prepared with Nextera DNA Flex or DNA Prep) (all Illumina). Lastly, sequencing on the NextSeq 1000, using the P2 600 cycle kit (for libraries prepared with DNA Prep) with onboard denaturation consisted of 10 ng inputs of each of the 372 donor libraries along with 8 positive and 4 negative controls pooled into a single tube. All library and sequencing methods were successfully validated using a blinded panel of samples from the 17th International HLA and Immunogenetics Workshop (IHIWS) as previously described^46,47^.

### 2.4 Bioinformatics and Statistical Analysis

All sequences generated by the MiSeq were analyzed using previously described parameters^35^. In brief, FASTQ files generated by MiSeq Reporter were analyzed in Omixon Target v1.8.0 or v1.9.3 (Omixon Biocomputing Kft, Budapest, Hungary) using IPD-IMGT/HLA database release 3.10-3.48, Omixon HLA Explore v2.0 using IPD-IMGT/HLA database release 3.51-3.56, and NGSengine v2.10.0-v3.30.0 with IPD-IMGT/HLA database releases 3.25-3.58 (GenDx, Utrecht, The Netherlands)^47,48^. For a subset of samples sequenced on the NextSeq 1000 instrument, DRAGEN BCL Convert version 4.2.7 (both Illumina) was used to generate FASTQ files for HLA genotyping analyses in NGSengine v3.2-3.30.0 with IPD-IMGT/HLA database references 3.56-3.58 (GenDX). Genotypes with initial novel or ambiguous allele combinations were periodically reanalyzed with each IPD-IMGT/HLA database release and updated accordingly. Manually selected alleles were reviewed in IGV using an in-house pipeline as described previously^35^. Population genetics analyses were performed using various software tools including Arlequin v3.5.2.2, PyPop v1.0.2 and Hapl-o-mat v1.3. Arlequin was used to calculate allele frequencies per locus at 2 and 3-field resolution. Total frequencies of rare alleles per locus were derived by the sum of all alleles with frequencies of <1.0%. Shannon and Simpson diversity indices were calculated per locus in each country using the diversity function from the vegan package (v2.5.7) in R^49^. Arlequin was used to calculate deviation from Hardy-Weinberg Equilibrium (HWE) using the Guo and Thompson implementation. The neutrality tests were performed using Slatkin’s implementation of Ewens-Watterson (EW) homozygosity test of neutrality in PyPop^50-52^. In Hapl-o-mat, haplotype frequencies were calculated for all pairwise combinations within class I and within class II loci, as well as each class I locus with DRB1, DPA1∼DPB1, DQA1∼DQB1, and DRB1∼DRB3/4/5. We also calculated haplotype frequencies for select multi-locus haplotypes: all class I, all class II, and all 11 genotyped loci. For ambiguous genotype calls the more common alleles were chosen as the HLA types for these analyses. DRB3, DRB4, and DRB5 genes that were not present in a given participant, due to the nature of the DRB haplotypes, were reported as the 00:00:00 allele. Individuals with repeated failed PCR amplifications were included in allele reporting with an allowable maximum of 1 locus missing/failed for inclusion. The sum allele frequencies for missing/failed or allele variants identified but not matching sequences in the IPD-IMGT/HLA database were reported as “Locus*00:00:00”. All sequences were submitted to GenBank and the IPD-IMGT/HLA databases and accession numbers are provided.

## 3.0 Results

### 3.1 Single-locus analyses in African populations

HLA genotyping was performed on 1023 individuals enrolled in various clinical studies from five African countries: Kenya, Uganda, Tanzania, Mozambique, and Nigeria (Fig. 1A). Overall, we identified 49 HLA-A, 72 B, 46 C, 30 DPA1, 65 DPB1, 24 DQA1, 28 DQB1, 42 DRB1, 7 DRB3, 4 DRB4, and 4 DRB5 alleles, for a total of 371 HLA alleles in these African populations (Fig. 1B-C, Tables 1-6, Supplementary Tables 1-3). The East African countries Kenya, Uganda, and Tanzania collectively had the highest number of total alleles for HLA-B (63), DPB1 (51) and A (45) loci (Tables 1-2, & 4). For Mozambique in the Southeast, the top three loci with the greatest number of total alleles were HLA-B (29), C (23) and DPB1 (25) loci (Tables 2-4). Nigeria in West Africa demonstrated its highest total alleles for HLA-B (36), A (30) and DRB1 (27) loci (Tables 1, 2, & 6). East African countries shared 158 alleles, which included 28 HLA-A, 29 B, 19 C, 10 DPA1, 20 DPB1, 12 DQA1, 15 DQB1, 18 DRB1, 4 DRB3, 2 DRB4, and 1 DRB5 allele (Tables 1-6, Supplementary Tables 1-3).

**Table 1.** HLA-A allele frequencies in Africa across East, Southeast, and West regions at 3-field resolution.

| Allele | Africa |  | East |  |  |  |  |  | Southeast |  | West |  |
| --- | --- | --- | --- | --- | --- | --- | --- | --- | --- | --- | --- | --- |
|  | Overall |  | Kenya |  | Uganda |  | Tanzania |  | Mozambique |  | Nigeria |  |
|  | Frequency | 2N=2046 | Frequency | 2N=750 | Frequency | 2N=676 | Frequency | 2N=178 | Frequency | 2N=164 | Frequency | 2N=278 |
| A*01:01:01 | 0.0484 | 99 | 0.0587 | 44 | 0.0621 | 42 | 0.0225 | 4 | 0.0305 | 5 | 0.0144 | 4 |
| A*01:01:83 | 0.0132 | 27 | 0.0227 | 17 | 0.0118 | 8 | 0.0056 | 1 | 0.0061 | 1 |  |  |
| A*01:02:01 | 0.0044 | 9 | 0.0040 | 3 | 0.0059 | 4 |  |  | 0.0061 | 1 | 0.0036 | 1 |
| A*01:03:01 | 0.0068 | 14 | 0.0107 | 8 | 0.0059 | 4 | 0.0112 | 2 |  |  |  |  |
| A*01:09:01 | 0.0020 | 4 | 0.0027 | 2 | 0.0015 | 1 | 0.0056 | 1 |  |  |  |  |
| A*01:23:01 | 0.0005 | 1 |  |  | 0.0015 | 1 |  |  |  |  |  |  |
| A*02:01:01 | 0.1310 | 268 | 0.1613 | 121 | 0.1213 | 82 | 0.1292 | 23 | 0.0793 | 13 | 0.1043 | 29 |
| A*02:02:01 | 0.0420 | 86 | 0.0347 | 26 | 0.0429 | 29 | 0.0337 | 6 | 0.0366 | 6 | 0.0684 | 19 |
| A*02:02:05 | 0.0015 | 3 | 0.0013 | 1 |  |  | 0.0112 | 2 |  |  |  |  |
| A*02:05:01 | 0.0293 | 60 | 0.0413 | 31 | 0.0104 | 7 | 0.0393 | 7 | 0.0671 | 11 | 0.0144 | 4 |
| A*02:14 | 0.0103 | 21 | 0.0080 | 6 | 0.0178 | 12 | 0.0169 | 3 |  |  |  |  |
| A*02:60:01 | 0.0010 | 2 |  |  |  |  |  |  |  |  | 0.0072 | 2 |
| A*02:85 | 0.0005 | 1 | 0.0013 | 1 |  |  |  |  |  |  |  |  |
| A*02:951 | 0.0005 | 1 |  |  | 0.0015 | 1 |  |  |  |  |  |  |
| A*03:01:01 | 0.0513 | 105 | 0.0480 | 36 | 0.0488 | 33 | 0.0562 | 10 | 0.0366 | 6 | 0.0719 | 20 |
| A*23:01:01 | 0.0640 | 131 | 0.0413 | 31 | 0.0621 | 42 | 0.0955 | 17 | 0.0488 | 8 | 0.1187 | 33 |
| A*23:17:01 | 0.0196 | 40 | 0.0147 | 11 | 0.0192 | 13 | 0.0225 | 4 | 0.0305 | 5 | 0.0252 | 7 |
| A*24:02:01 | 0.0186 | 38 | 0.0320 | 24 | 0.0104 | 7 | 0.0056 | 1 | 0.0183 | 3 | 0.0108 | 3 |
| A*26:01:01 | 0.0059 | 12 | 0.0067 | 5 | 0.0059 | 4 | 0.0056 | 1 | 0.0122 | 2 |  |  |
| A*26:12:01 | 0.0064 | 13 | 0.0093 | 7 | 0.0059 | 4 |  |  |  |  | 0.0072 | 2 |
| A*26:30 | 0.0020 | 4 | 0.0053 | 4 |  |  |  |  |  |  |  |  |
| A*29:01:01 | 0.0122 | 25 | 0.0147 | 11 | 0.0192 | 13 | 0.0056 | 1 |  |  |  |  |
| A*29:02:01 | 0.0494 | 101 | 0.0427 | 32 | 0.0533 | 36 | 0.0393 | 7 | 0.1037 | 17 | 0.0324 | 9 |
| A*29:11 | 0.0005 | 1 |  |  |  |  |  |  | 0.0061 | 1 |  |  |
| A*29:15 | 0.0020 | 4 | 0.0053 | 4 |  |  |  |  |  |  |  |  |
| A*30:01:01 | 0.0645 | 132 | 0.0560 | 42 | 0.0592 | 40 | 0.0787 | 14 | 0.0915 | 15 | 0.0755 | 21 |
| A*30:02:01 | 0.0694 | 142 | 0.0507 | 38 | 0.0902 | 61 | 0.0787 | 14 | 0.1159 | 19 | 0.0360 | 10 |
| A*30:04:01 | 0.0112 | 23 | 0.0147 | 11 | 0.0118 | 8 | 0.0056 | 1 | 0.0122 | 2 | 0.0036 | 1 |
| A*30:09:01 | 0.0010 | 2 | 0.0013 | 1 | 0.0015 | 1 |  |  |  |  |  |  |
| A*31:01:02 | 0.0039 | 8 | 0.0013 | 1 | 0.0074 | 5 |  |  |  |  | 0.0072 | 2 |
| A*31:04:01 | 0.0059 | 12 | 0.0107 | 8 | 0.0059 | 4 |  |  |  |  |  |  |
| A*32:01:01 | 0.0122 | 25 | 0.0093 | 7 | 0.0118 | 8 |  |  | 0.0061 | 1 | 0.0324 | 9 |
| A*32:106:01 | 0.0005 | 1 | 0.0013 | 1 |  |  |  |  |  |  |  |  |
| A*33:01:01 | 0.0059 | 12 | 0.0067 | 5 | 0.0030 | 2 | 0.0056 | 1 |  |  | 0.0144 | 4 |
| A*33:03:01 | 0.0196 | 40 | 0.0147 | 11 | 0.0118 | 8 | 0.0281 | 5 | 0.0122 | 2 | 0.0504 | 14 |
| A*34:02:01 | 0.0279 | 57 | 0.0293 | 22 | 0.0178 | 12 | 0.0169 | 3 | 0.0244 | 4 | 0.0576 | 16 |
| A*36:01:01 | 0.0347 | 71 | 0.0147 | 11 | 0.0281 | 19 | 0.1067 | 19 | 0.0244 | 4 | 0.0648 | 18 |
| A*66:01:01 | 0.0440 | 90 | 0.0413 | 31 | 0.0577 | 39 | 0.0225 | 4 | 0.0610 | 10 | 0.0216 | 6 |
| A*66:02:01 | 0.0068 | 14 | 0.0040 | 3 | 0.0044 | 3 | 0.0056 | 1 |  |  | 0.0252 | 7 |
| A*66:03:01 | 0.0020 | 4 | 0.0013 | 1 | 0.0030 | 2 |  |  |  |  | 0.0036 | 1 |
| A*68:01:01 | 0.0152 | 31 | 0.0133 | 10 | 0.0148 | 10 | 0.0169 | 3 | 0.0366 | 6 | 0.0072 | 2 |
| A*68:02:01 | 0.0782 | 160 | 0.1000 | 75 | 0.0666 | 45 | 0.0843 | 15 | 0.0610 | 10 | 0.0540 | 15 |
| A*68:278 | 0.0010 | 2 | 0.0027 | 2 |  |  |  |  |  |  |  |  |
| A*74:01:01 | 0.0596 | 122 | 0.0480 | 36 | 0.0828 | 56 | 0.0393 | 7 | 0.0549 | 9 | 0.0504 | 14 |
| A*74:03:01 | 0.0073 | 15 | 0.0053 | 4 | 0.0148 | 10 | 0.0056 | 1 |  |  |  |  |
| A*74:09 | 0.0005 | 1 |  |  |  |  |  |  |  |  | 0.0036 | 1 |
| A*80:01:01 | 0.0039 | 8 | 0.0027 | 2 |  |  |  |  | 0.0183 | 3 | 0.0108 | 3 |
| A*00:00:00 | 0.0020 | 4 | 0.0040 | 3 |  |  |  |  |  |  | 0.0036 | 1 |

Overall allele frequencies at 3-field resolution for HLA-A locus were calculated per population.
The 2N values reflect total number of alleles found per HLA-A allele. "A\*00:00:00" reflects missing data which includes novel alleles awaiting further or final confirmation.

**Table 2.** HLA-B allele frequencies in Africa across East, Southeast, and West regions at 3-field resolution.

| Allele | Africa |  | East |  | East |  | East |  | Southeast |  | West |  |
| --- | --- | --- | --- | --- | --- | --- | --- | --- | --- | --- | --- | --- |
|  | Overall |  | Kenya |  | Uganda |  | Tanzania |  | Mozambique |  | Nigeria |  |
|  | Frequency | 2N=2046 | Frequency | 2N=750 | Frequency | 2N=676 | Frequency | 2N=178 | Frequency | 2N=164 | Frequency | 2N=278 |
| B*07:02:01 | 0.0494 | 101 | 0.0400 | 30 | 0.0488 | 33 | 0.0787 | 14 | 0.0122 | 2 | 0.0791 | 22 |
| B*07:05:01 | 0.0049 | 10 | 0.0107 | 8 | 0.0030 | 2 |  |  |  |  |  |  |
| B*07:06:01 | 0.0005 | 1 | 0.0013 | 1 |  |  |  |  |  |  |  |  |
| B*08:01:01 | 0.0318 | 65 | 0.0320 | 24 | 0.0370 | 25 | 0.0337 | 6 | 0.0427 | 7 | 0.0108 | 3 |
| B*13:02:01 | 0.0127 | 26 | 0.0093 | 7 | 0.0163 | 11 | 0.0281 | 5 | 0.0122 | 2 | 0.0036 | 1 |
| B*13:03 | 0.0049 | 10 | 0.0107 | 8 | 0.0015 | 1 |  |  | 0.0061 | 1 |  |  |
| B*14:01:01 | 0.0054 | 11 | 0.0027 | 2 | 0.0044 | 3 | 0.0056 | 1 | 0.0183 | 3 | 0.0072 | 2 |
| B*14:02:01 | 0.0225 | 46 | 0.0173 | 13 | 0.0340 | 23 | 0.0281 | 5 | 0.0244 | 4 | 0.0036 | 1 |
| B*14:03 | 0.0015 | 3 |  |  |  |  |  |  |  |  | 0.0108 | 3 |
| B*15:01:01 | 0.0005 | 1 |  |  |  |  |  |  | 0.0061 | 1 |  |  |
| B*15:03:01 | 0.0616 | 126 | 0.0680 | 51 | 0.0754 | 51 | 0.0449 | 8 | 0.0427 | 7 | 0.0324 | 9 |
| B*15:10:01 | 0.0469 | 96 | 0.0267 | 20 | 0.0459 | 31 | 0.0955 | 17 | 0.0671 | 11 | 0.0612 | 17 |
| B*15:16:01 | 0.0073 | 15 | 0.0054 | 4 | 0.0074 | 5 | 0.0056 | 1 |  |  | 0.0180 | 5 |
| B*15:17:01 | 0.0015 | 3 | 0.0027 | 2 |  |  |  |  |  |  | 0.0036 | 1 |
| B*15:18:01 | 0.0015 | 3 |  |  | 0.0015 | 1 |  |  |  |  | 0.0072 | 2 |
| B*15:220:01 | 0.0103 | 21 | 0.0187 | 14 | 0.0089 | 6 |  |  | 0.0061 | 1 |  |  |
| B*15:31:01 | 0.0015 | 3 |  |  | 0.0030 | 2 | 0.0056 | 1 |  |  |  |  |
| B*15:37 | 0.0010 | 2 | 0.0013 | 1 | 0.0015 | 1 |  |  |  |  |  |  |
| B*15:64:02 | 0.0005 | 1 | 0.0013 | 1 |  |  |  |  |  |  |  |  |
| B*15:67 | 0.0034 | 7 | 0.0080 | 6 | 0.0015 | 1 |  |  |  |  |  |  |
| B*18:01:01 | 0.0327 | 67 | 0.0347 | 26 | 0.0296 | 20 | 0.0225 | 4 | 0.0671 | 11 | 0.0216 | 6 |
| B*18:03:01 | 0.0049 | 10 | 0.0027 | 2 | 0.0089 | 6 | 0.0056 | 1 | 0.0061 | 1 |  |  |
| B*27:03 | 0.0098 | 20 | 0.0160 | 12 | 0.0059 | 4 |  |  |  |  | 0.0144 | 4 |
| B*27:05:02 | 0.0005 | 1 |  |  | 0.0015 | 1 |  |  |  |  |  |  |
| B*35:01:01 | 0.0308 | 63 | 0.0333 | 25 | 0.0133 | 9 | 0.0169 | 3 | 0.0183 | 3 | 0.0827 | 23 |
| B*35:02:01 | 0.0068 | 14 | 0.0160 | 12 | 0.0015 | 1 | 0.0056 | 1 |  |  |  |  |
| B*35:03:01 | 0.0005 | 1 |  |  | 0.0015 | 1 |  |  |  |  |  |  |
| B*35:189 | 0.0005 | 1 | 0.0013 | 1 |  |  |  |  |  |  |  |  |
| B*37:01:01 | 0.0034 | 7 | 0.0054 | 4 | 0.0030 | 2 |  |  |  |  | 0.0036 | 1 |
| B*38:01:01 | 0.0005 | 1 |  |  |  |  | 0.0056 | 1 |  |  |  |  |
| B*39:01:01 | 0.0005 | 1 | 0.0013 | 1 |  |  |  |  |  |  |  |  |
| B*39:06:02 | 0.0005 | 1 | 0.0013 | 1 |  |  |  |  |  |  |  |  |
| B*39:10:01 | 0.0156 | 32 | 0.0160 | 12 | 0.0148 | 10 | 0.0169 | 3 | 0.0061 | 1 | 0.0216 | 6 |
| B*39:24:01 | 0.0034 | 7 | 0.0067 | 5 | 0.0015 | 1 |  |  |  |  | 0.0036 | 1 |
| B*40:02:01 | 0.0005 | 1 |  |  |  |  |  |  |  |  | 0.0036 | 1 |
| B*40:12:01 | 0.0049 | 10 | 0.0080 | 6 | 0.0059 | 4 |  |  |  |  |  |  |
| B*40:16:01 | 0.0024 | 5 | 0.0040 | 3 |  |  |  |  | 0.0122 | 2 |  |  |
| B*41:01:01 | 0.0132 | 27 | 0.0267 | 20 | 0.0089 | 6 |  |  | 0.0061 | 1 |  |  |
| B*41:02:01 | 0.0054 | 11 | 0.0093 | 7 | 0.0059 | 4 |  |  |  |  |  |  |
| B*42:01:01 | 0.0621 | 127 | 0.0627 | 47 | 0.0681 | 46 | 0.0562 | 10 | 0.0488 | 8 | 0.0576 | 16 |
| B*42:02:01 | 0.0044 | 9 | 0.0013 | 1 | 0.0030 | 2 | 0.0056 | 1 | 0.0244 | 4 | 0.0036 | 1 |
| B*44:03:01 | 0.0288 | 59 | 0.0307 | 23 | 0.0133 | 9 | 0.0169 | 3 | 0.0671 | 11 | 0.0468 | 13 |
| B*44:03:02 | 0.0093 | 19 | 0.0027 | 2 | 0.0015 | 1 |  |  | 0.0732 | 12 | 0.0144 | 4 |
| B*44:07 | 0.0010 | 2 |  |  |  |  |  |  |  |  | 0.0072 | 2 |
| B*44:15:01 | 0.0073 | 15 | 0.0067 | 5 | 0.0133 | 9 | 0.0056 | 1 |  |  |  |  |
| B*45:01:01 | 0.0714 | 146 | 0.0693 | 52 | 0.0888 | 60 | 0.1124 | 20 | 0.0305 | 5 | 0.0324 | 9 |
| B*45:07 | 0.0024 | 5 | 0.0027 | 2 | 0.0015 | 1 | 0.0056 | 1 | 0.0061 | 1 |  |  |
| B*47:01:01 | 0.0010 | 2 | 0.0013 | 1 |  |  | 0.0056 | 1 |  |  |  |  |
| B*47:03 | 0.0054 | 11 | 0.0107 | 8 | 0.0044 | 3 |  |  |  |  |  |  |
| B*48:05:01 | 0.0015 | 3 | 0.0013 | 1 | 0.0015 | 1 | 0.0056 | 1 |  |  |  |  |
| B*49:01:01 | 0.0459 | 94 | 0.0533 | 40 | 0.0547 | 37 | 0.0112 | 2 | 0.0488 | 8 | 0.0252 | 7 |
| B*50:01:01 | 0.0015 | 3 | 0.0013 | 1 | 0.0015 | 1 | 0.0056 | 1 |  |  |  |  |
| B*51:01:01 | 0.0205 | 42 | 0.0333 | 25 | 0.0104 | 7 | 0.0225 | 4 |  |  | 0.0216 | 6 |
| B*52:01:02 | 0.0029 | 6 |  |  |  |  |  |  |  |  | 0.0216 | 6 |
| B*53:01:01 | 0.0953 | 195 | 0.0547 | 41 | 0.0976 | 66 | 0.1461 | 26 | 0.0854 | 14 | 0.1727 | 48 |
| B*53:08:01 | 0.0005 | 1 |  |  | 0.0015 | 1 |  |  |  |  |  |  |
| B*53:37 | 0.0005 | 1 |  |  |  |  |  |  |  |  | 0.0036 | 1 |
| B*56:01:01 | 0.0020 | 4 | 0.0040 | 3 | 0.0015 | 1 |  |  |  |  |  |  |
| B*57:01:01 | 0.0044 | 9 | 0.0120 | 9 |  |  |  |  |  |  |  |  |
| B*57:02:01 | 0.0068 | 14 | 0.0093 | 7 | 0.0089 | 6 |  |  | 0.0061 | 1 |  |  |
| B*57:03:01 | 0.0274 | 56 | 0.0253 | 19 | 0.0281 | 19 | 0.0169 | 3 | 0.0183 | 3 | 0.0432 | 12 |
| B*57:05 | 0.0005 | 1 |  |  |  |  |  |  |  |  | 0.0036 | 1 |
| B*58:01:01 | 0.0616 | 126 | 0.0600 | 45 | 0.0533 | 36 | 0.0506 | 9 | 0.1037 | 17 | 0.0684 | 19 |
| B*58:02:01 | 0.0816 | 167 | 0.0680 | 51 | 0.1021 | 69 | 0.0955 | 17 | 0.0976 | 16 | 0.0504 | 14 |
| B*58:64 | 0.0005 | 1 | 0.0013 | 1 |  |  |  |  |  |  |  |  |
| B*73:01:01 | 0.0020 | 4 | 0.0027 | 2 | 0.0015 | 1 | 0.0056 | 1 |  |  |  |  |
| B*78:01:01 | 0.0015 | 3 |  |  |  |  |  |  |  |  | 0.0108 | 3 |
| B*81:01:01 | 0.0376 | 77 | 0.0387 | 29 | 0.0429 | 29 | 0.0337 | 6 | 0.0366 | 6 | 0.0252 | 7 |
| B*82:01:01 | 0.0005 | 1 |  |  |  |  |  |  |  |  | 0.0036 | 1 |
| B*82:02:01 | 0.0039 | 8 | 0.0040 | 3 | 0.0074 | 5 |  |  |  |  |  |  |
| B*00:00:00 | 0.0020 | 4 | 0.0040 | 3 | 0.0015 | 1 |  |  |  |  |  |  |

Overall allele frequencies at 3-field resolution for HLA-B locus were calculated per population.
The 2N values reflect total number of alleles found per HLA-B allele. "B\*00:00:00" reflects missing data which includes novel alleles awaiting further or final confirmation.

**Table 3.**
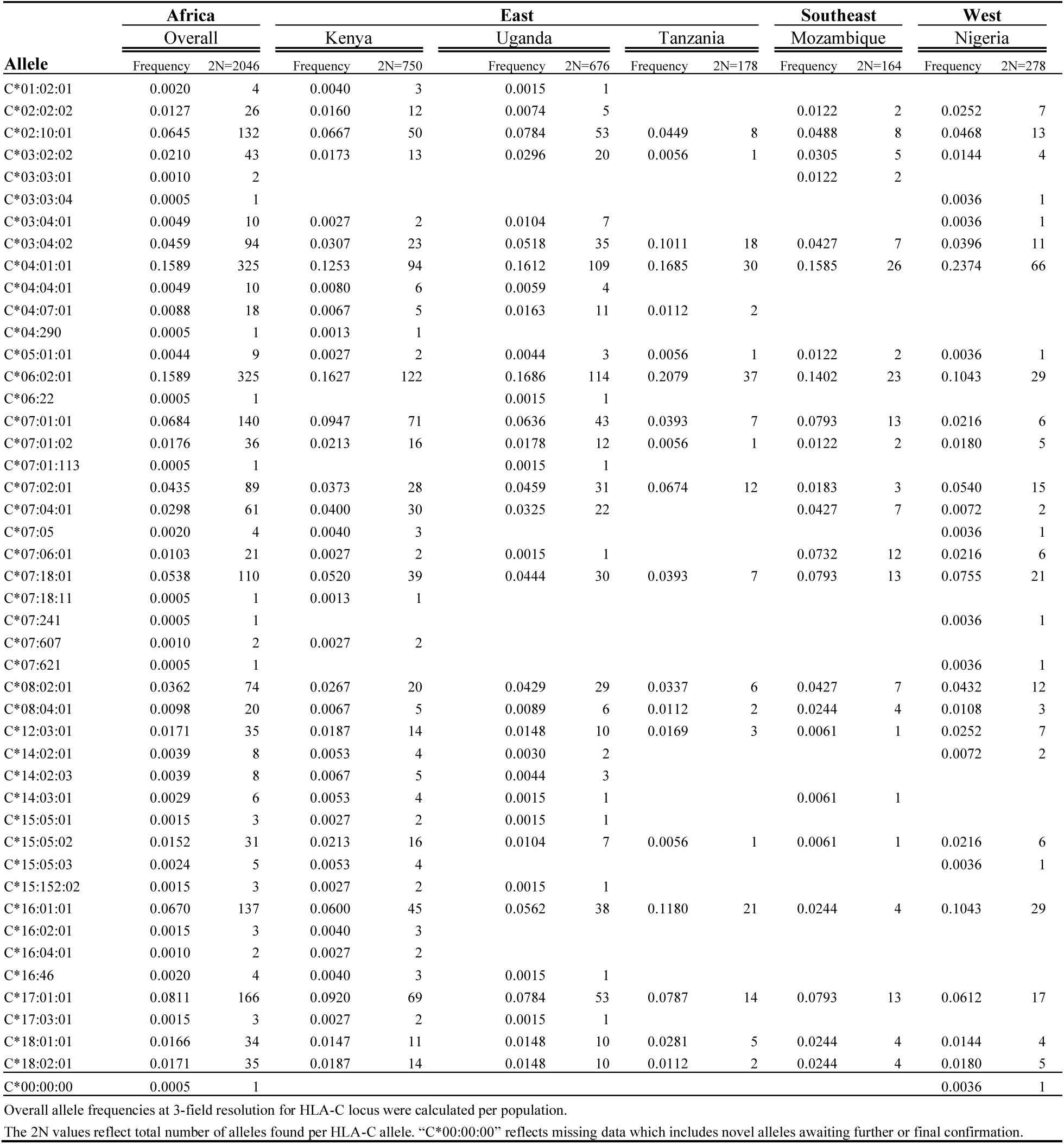
HLA-C allele frequencies in Africa across East, Southeast, and West regions at 3-field resolution.

**Table 4.**
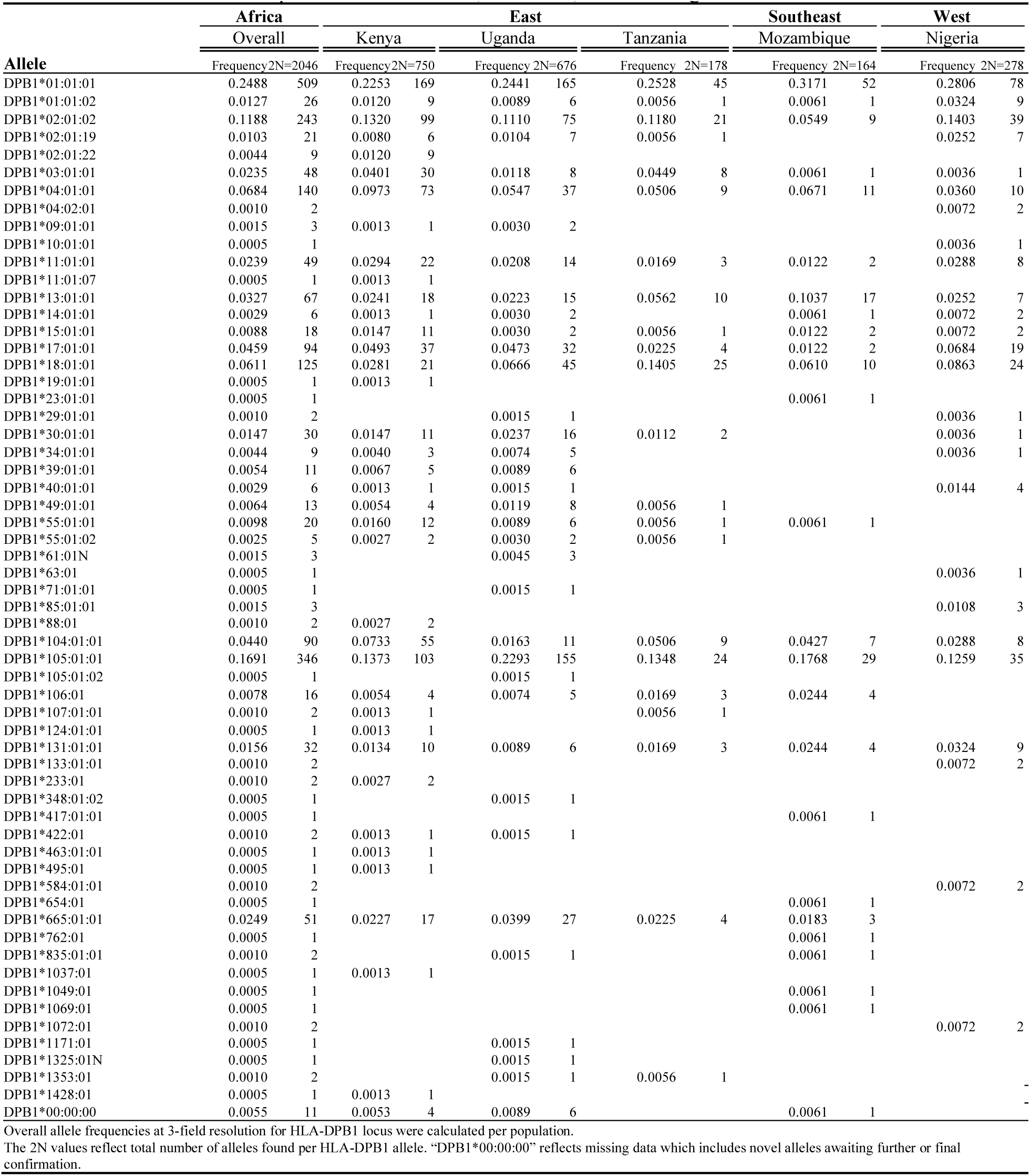
HLA-DPB1 allele frequencies in Africa East, Southeast, and West regions at 3-field resolution.

**Table 5.**
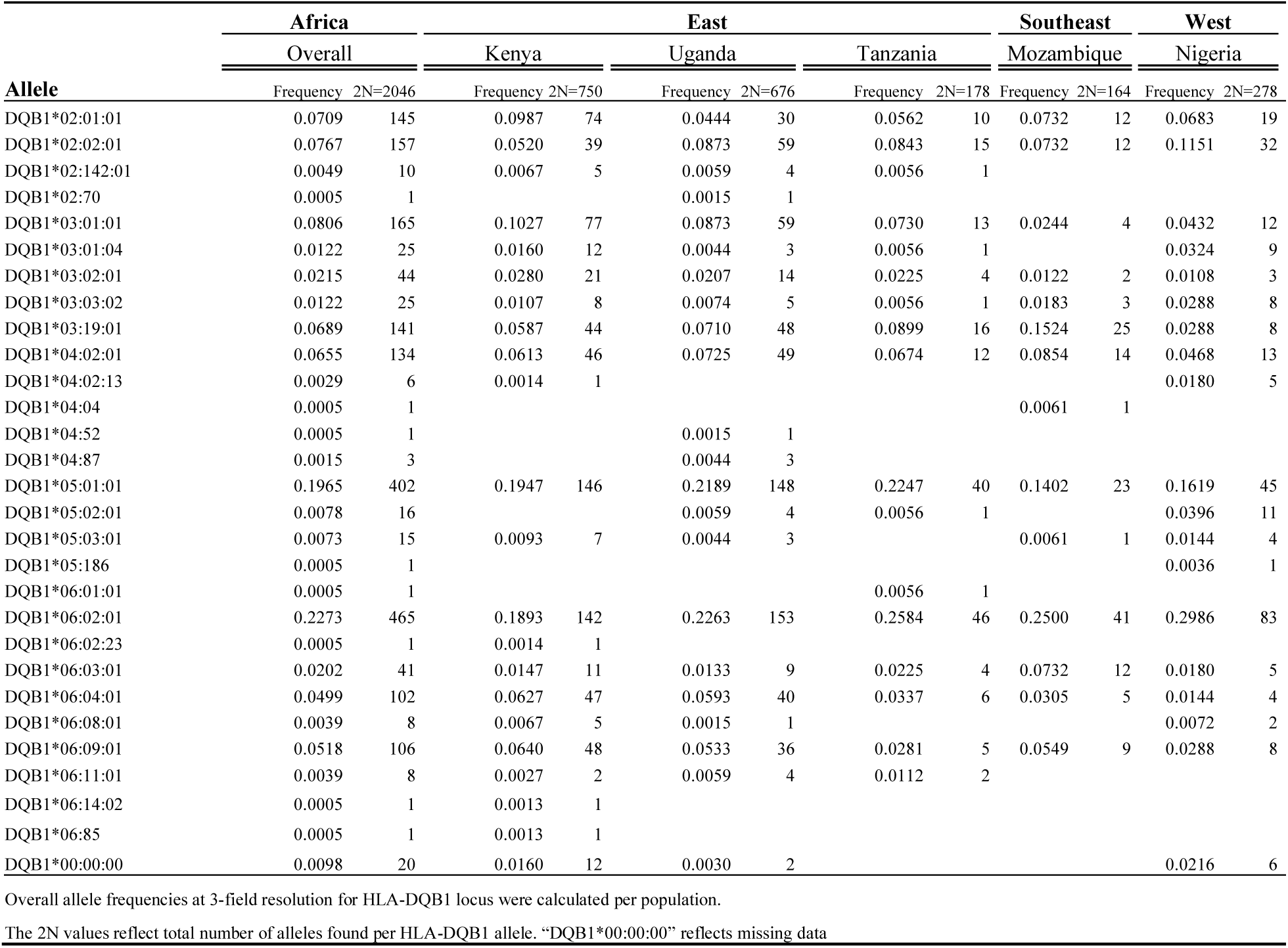
HLA-DQB1 allele frequencies in Africa across East, Southeast, and West regions at 3-field resolution.

**Table 6.** HLA-DRB1 allele frequencies in Africa across East, Southeast, and West regions at 3-field resolution.

| Allele | Africa |  | East |  |  |  |  |  | Southeast |  | West |  |
| --- | --- | --- | --- | --- | --- | --- | --- | --- | --- | --- | --- | --- |
|  | Overall |  | Kenya |  | Uganda |  | Tanzania |  | Mozambique |  | Nigeria |  |
|  | Frequency | 2N=2046 | Frequency | 2N=750 | Frequency | 2N=676 | Frequency | 2N=178 | Frequency | 2N=164 | Frequency | 2N=278 |
| DRB1*01:01:01 | 0.0117 | 24 | 0.0107 | 8 | 0.0133 | 9 | 0.0225 | 4 | 0.0122 | 2 | 0.0036 | 1 |
| DRB1*01:02:01 | 0.0611 | 125 | 0.0693 | 52 | 0.0636 | 43 | 0.0730 | 13 | 0.0305 | 5 | 0.0432 | 12 |
| DRB1*03:01:01 | 0.0718 | 147 | 0.1080 | 81 | 0.0399 | 27 | 0.0506 | 9 | 0.0671 | 11 | 0.0683 | 19 |
| DRB1*03:01:02 | 0.0005 | 1 |  |  |  |  |  |  |  |  | 0.0036 | 1 |
| DRB1*03:02:01 | 0.0719 | 147 | 0.0720 | 54 | 0.0740 | 50 | 0.0730 | 13 | 0.0915 | 15 | 0.0540 | 15 |
| DRB1*04:01:01 | 0.0073 | 15 | 0.0147 | 11 | 0.0030 | 2 |  |  | 0.0061 | 1 | 0.0036 | 1 |
| DRB1*04:04:01 | 0.0015 | 3 | 0.0040 | 3 |  |  |  |  |  |  |  |  |
| DRB1*04:05:01 | 0.0191 | 39 | 0.0267 | 20 | 0.0178 | 12 | 0.0225 | 4 | 0.0061 | 1 | 0.0072 | 2 |
| DRB1*04:10:01 | 0.0005 | 1 |  |  | 0.0015 | 1 |  |  |  |  |  |  |
| DRB1*07:01:01 | 0.0547 | 112 | 0.0507 | 38 | 0.0562 | 38 | 0.0506 | 9 | 0.0549 | 9 | 0.0647 | 18 |
| DRB1*08:04:01 | 0.0435 | 89 | 0.0440 | 33 | 0.0355 | 24 | 0.0225 | 4 | 0.0183 | 3 | 0.0899 | 25 |
| DRB1*08:06:01 | 0.0005 | 1 |  |  |  |  |  |  |  |  | 0.0036 | 1 |
| DRB1*08:08 | 0.0015 | 3 |  |  | 0.0044 | 3 |  |  |  |  |  |  |
| DRB1*09:01:02 | 0.0181 | 37 | 0.0040 | 3 | 0.0222 | 15 | 0.0393 | 7 | 0.0183 | 3 | 0.0324 | 9 |
| DRB1*10:01:01 | 0.0391 | 80 | 0.0413 | 31 | 0.0370 | 25 | 0.0506 | 9 | 0.0427 | 7 | 0.0288 | 8 |
| DRB1*11:01:01 | 0.0024 | 5 | 0.0013 | 1 | 0.0030 | 2 |  |  |  |  | 0.0072 | 2 |
| DRB1*11:01:02 | 0.1422 | 291 | 0.1387 | 104 | 0.1553 | 105 | 0.1685 | 30 | 0.1707 | 28 | 0.0863 | 24 |
| DRB1*11:02:01 | 0.0596 | 122 | 0.0520 | 39 | 0.0754 | 51 | 0.0506 | 9 | 0.0915 | 15 | 0.0288 | 8 |
| DRB1*11:04:01 | 0.0059 | 12 | 0.0093 | 7 | 0.0030 | 2 | 0.0169 | 3 |  |  |  |  |
| DRB1*11:04:02 | 0.0010 | 2 |  |  |  |  |  |  |  |  | 0.0072 | 2 |
| DRB1*11:08:02 | 0.0005 | 1 |  |  |  |  |  |  |  |  | 0.0036 | 1 |
| DRB1*11:10:01 | 0.0010 | 2 |  |  |  |  |  |  |  |  | 0.0072 | 2 |
| DRB1*11:14:02 | 0.0010 | 2 |  |  |  |  | 0.0056 | 1 | 0.0061 | 1 |  |  |
| DRB1*12:01:01 | 0.0303 | 62 | 0.0160 | 12 | 0.0459 | 31 | 0.0337 | 6 | 0.0183 | 3 | 0.0360 | 10 |
| DRB1*12:02:02 | 0.0015 | 3 |  |  | 0.0015 | 1 |  |  | 0.0122 | 2 |  |  |
| DRB1*13:01:01 | 0.0616 | 126 | 0.0493 | 37 | 0.0429 | 29 | 0.0674 | 12 | 0.1281 | 21 | 0.0971 | 27 |
| DRB1*13:01:03 | 0.0005 | 1 |  |  |  |  |  |  |  |  | 0.0036 | 1 |
| DRB1*13:02:01 | 0.0924 | 189 | 0.1147 | 86 | 0.1095 | 74 | 0.0449 | 8 | 0.0793 | 13 | 0.0288 | 8 |
| DRB1*13:03:01 | 0.0283 | 58 | 0.0320 | 24 | 0.0266 | 18 | 0.0169 | 3 |  |  | 0.0468 | 13 |
| DRB1*14:01:02 | 0.0005 | 1 |  |  | 0.0015 | 1 |  |  |  |  |  |  |
| DRB1*14:54:01 | 0.0108 | 22 | 0.0107 | 8 | 0.0104 | 7 | 0.0112 | 2 | 0.0061 | 1 | 0.0144 | 4 |
| DRB1*14:243 | 0.0010 | 2 |  |  |  |  |  |  | 0.0061 | 1 | 0.0036 | 1 |
| DRB1*15:01:01 | 0.0015 | 3 |  |  | 0.0030 | 2 |  |  | 0.0061 | 1 |  |  |
| DRB1*15:02:01 | 0.0005 | 1 |  |  |  |  | 0.0056 | 1 |  |  |  |  |
| DRB1*15:03:01 | 0.1486 | 304 | 0.1280 | 96 | 0.1465 | 99 | 0.1629 | 29 | 0.1220 | 20 | 0.2158 | 60 |
| DRB1*16:02:01 | 0.0029 | 6 |  |  | 0.0030 | 2 | 0.0056 | 1 |  |  | 0.0108 | 3 |
| DRB1*00:00:00 | 0.0035 | 7 | 0.0026 | 2 | 0.0045 | 3 | 0.0056 | 1 | 0.0061 | 1 |  |  |

Overall allele frequencies at 3-field resolution for HLA-DRB1 locus were calculated per population.
The 2N values reflect total number of alleles found per HLA-DRB1 allele. "DRB1\*00:00:00" reflects missing data which includes novel alleles awaiting further or final confirmation.

Single-locus frequencies were computed for all populations. The most frequent HLA class I alleles (per locus) with the most shared frequencies in East Africa included: A*02:01:01 (14.1%), B*58:02:01 (8.5%), and C*06:02:01 (17.0%) (Tables 1-3, Fig. 2A-C). In Mozambique, the most frequent HLA alleles included: A*30:02:01 (11.6%), B*58:01:01 (10.4%), and C*04:01:01 (15.9%) (Tables 1-3, Fig. 2A-C). In Nigeria, HLA class I alleles with the greatest frequencies were A*23:01:01 (11.9%), B*53:01:01 (17.3%), and C*04:01:01 (23.7%) (Tables 1-3, Fig. 2A-C).

**Figure 2.**
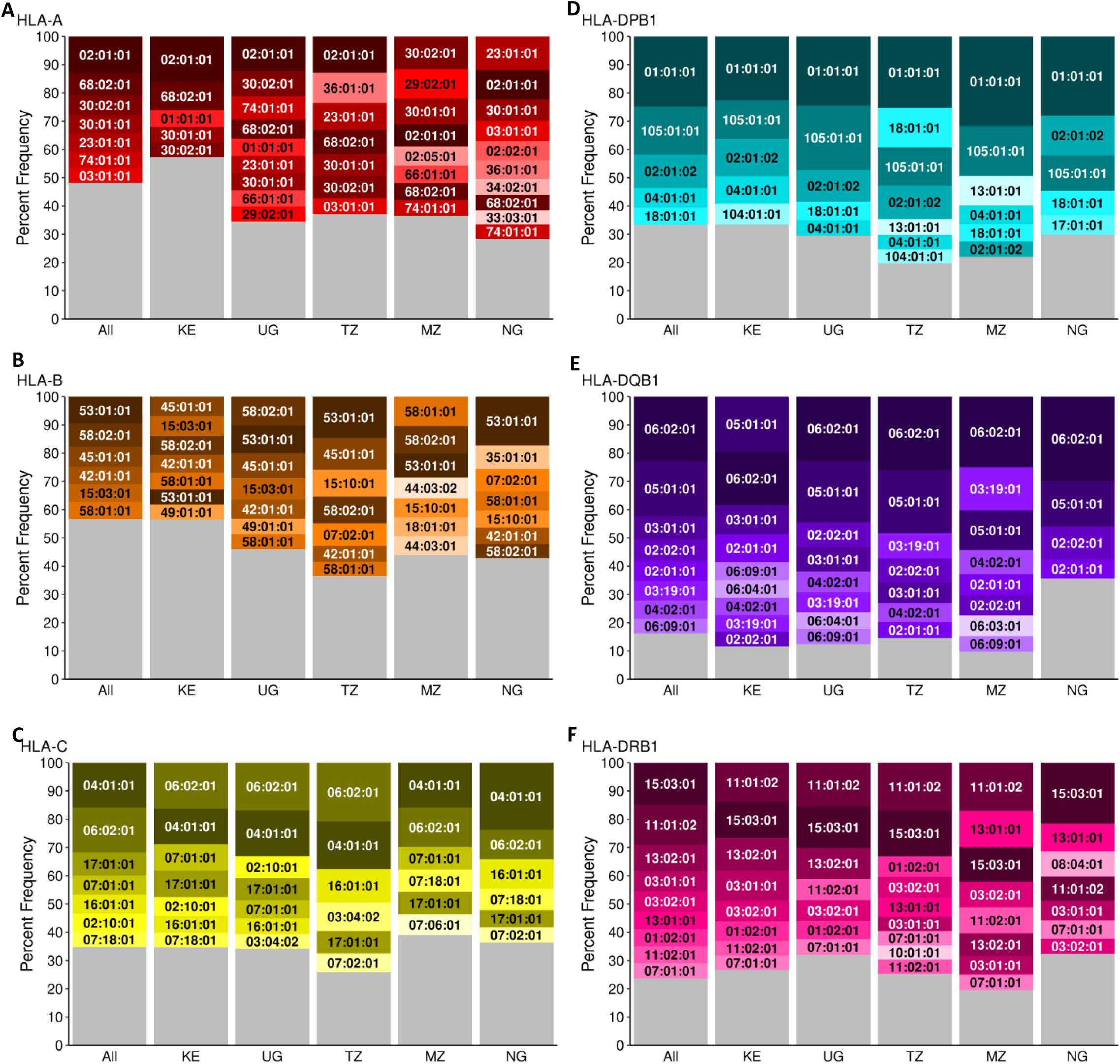
Common HLA class I and II frequencies of 1023 individuals in African countries (All), Kenya (KE), Uganda (UG), Tanzania (TZ), Mozambique (MZ), and Nigeria (NG). **A.** HLA-A, **B.** HLA-B, **C.** HLA-C, **D.** HLA-DPB1, **E.** HLA-DQB1, and **F.** HLA-DRB1. Alleles with percent frequencies >5% are represented in color over the sum of all alleles with <5% frequencies, shown in grey. Shared alleles are highlighted with the same color per locus across countries.

The most frequent HLA alleles for each class II locus in East Africa were DPA1*01:03:01 (37.6%), DPB1*01:01:01 (23.6%), DQA1*01:02:01 (31.3%), DQB1*06:02:01 (21.3%), DRB1*11:01:02 (14.9%), DRB3*02:02:01 (31.6%), DRB4*01:03:01 (7.2%), and DRB5*01:01:01 (14.2%) (Fig. 2D-F, Tables 4-6, Supplementary Fig. 1, Supplementary Tables 1-3). Similarly, in Mozambique, with the exception of DRB4, the most frequent alleles were identical to those in the other Eastern countries: DPA1*01:03:01 (25%), DPB1*01:01:01 (31.7%), DQA1*01:02:01 (32.9%), DQB1*06:02:01 (25%), DRB1*11:01:02 (17.1%), DRB3*02:02:01 (34.8%), for DRB4*01:01:01 (4.9%), and DRB5*01:01:01 (11.6%) (Fig. 2D-F, Tables 4-6, Supplementary Fig. 1, Supplementary Tables 1-3). In Nigeria, most of the top class II alleles were the same as those identified in East Africa (DPA1*01:03:01 (35.3%), DPB1*01:01:01 (28.1%), DQA1*01:02:01 (35.6%), DQB1*06:02:01 (29.9%), DRB3*02:02:01 (29.5%), and DRB5*01:01:01 (20.5%)), with the exception of DRB1*15:03:01 (21.6%) and DRB4*01:01:01 (6.5%) (Fig. 2D-F, Tables 4-6, Supplementary Fig. 1, Supplementary Tables 1-3).

Allele diversity analyses of HLA loci using the Shannon diversity index (richness and evenness of alleles) revealed the greatest diversity in African countries with larger sample sizes. Kenya exhibited the highest diversity at most loci, however the expected effect in Shannon diversity index with increased sample size deviated in Mozambique (DPA1) and Nigeria (DRB1) (Supplementary Table 4). Furthermore, the weighted allelic diversity of HLA loci determined by the Simpson diversity index indicated the highest abundance of diversity in Kenya with deviation from this pattern in Uganda (A) and Mozambique (DPA1) irrespective of sample size (Supplementary Table 4). Significant deviations from Hardy-Weinberg Equilibrium (HWE) were observed in the HLA-DQB1 and DRB3 loci in the overall African population (n=1023) (Supplementary Table 4). Population-specific deviations from HWE were observed in Kenya (DPB1, DQB1, and DRB3) and Tanzania (DRB3 and DRB4) (Supplementary Table 4). The Ewens-Watterson (EW) homozygosity test of neutrality showed a significant deviation from expected in the overall African population for HLA-A (p <0.025) (Supplementary Table 4). Population-specific EW significance was observed in Kenya (B and DRB1), Uganda (A), Mozambique (A, B, and DRB3), and Nigeria (A) (Supplementary Table 4).

### 3.2. Multi-locus analyses in African populations

Population-specific multi-locus haplotypes of HLA class I and II have been reported for various global populations. Studies have demonstrated the complexity of the strong linkage disequilibrium within the human MHC region with long range nonrandom linkage of HLA alleles. Here we calculated haplotype pairs within class I and class II in the overall African population and for each population. In the total African study, 30 haplotype pairs were found at >5% frequency (Table 7). Population-specific haplotypes with frequencies of >5% (but <5% in the overall African cohort) were found in Tanzania (27), Mozambique (19), Nigeria (15), Uganda (14), and Kenya (5) (Supplementary Table 5). Among the frequent pairwise haplotypes in the overall African cohort, there were fewer class I haplotypes (4) than those found for class II (26). The class I haplotypes were B*58:02:01∼C*06:02:01 (7.9%), B*53:01:01∼C*04:01:01 (7.5%), B*42:01:01∼C*17:01:01 (6.1%), and B*15:03:01∼C*02:10:01 (5.8%) (Table 7). Of the 26 common pairwise haplotypes in class II, 7 were found at greater than 10% frequency in the overall African cohort (Table 7). The most frequent class II haplotype was DQA1*01:02:01∼DQB1*06:02:01 (20.7%), which was also the most frequent in all countries (Table 7). The other common class II haplotypes that were also >10% in all populations were DPA1*02:02:02∼DPB1*01:01:01 (16.7%), DRB1*15:03:01∼DRB5*01:01:01 (14.8%), and DQB1*06:02:01∼DRB1*15:03:01 (14.4%) (Table 7).

**Table 7.** Pairwise haplotype frequencies (>5%) in Africa across East, Southeast, and West countries at 3-field resolution.

| Haplotype | Africa | East |  |  | Southeast | West |
| --- | --- | --- | --- | --- | --- | --- |
|  | Overall | Kenya | Uganda | Tanzania | Mozambique | Nigeria |
|  | 2N=2046 | 2N=750 | 2N=676 | 2N=178 | 2N=164 | 2N=278 |
| DQA1*01:02:01~DQB1*06:02:01 | 0.2073 | 0.1622 | 0.2085 | 0.2416 | 0.2277 | 0.2938 |
| DPA1*02:02:02~DPB1*01:01:01 | 0.1665 | 0.1591 | 0.1573 | 0.1854 | 0.2256 | 0.1619 |
| DRB1*15:03:01~DRB5*01:01:01 | 0.1480 | 0.1287 | 0.1458 | 0.1685 | 0.1188 | 0.2096 |
| DQB1*06:02:01~DRB1*15:03:01 | 0.1444 | 0.1214 | 0.1454 | 0.1517 | 0.1220 | 0.2116 |
| DPA1*03:01:01~DPB1*105:01:01 | 0.1386 | 0.1082 | 0.1958 | 0.0955 | 0.1707 | 0.0935 |
| DRB1*11:01:02~DRB3*02:02:01 | 0.1261 | 0.1227 | 0.1367 | 0.1532 | 0.1688 | 0.0642 |
| DPA1*01:03:01~DPB1*02:01:02 | 0.1185 | 0.1309 | 0.1113 | 0.1180 | 0.0549 | 0.1403 |
| DRB1*13:02:01~DRB3*03:01:01 | 0.0926 | 0.1138 | 0.1101 | 0.0449 | 0.0813 | 0.0294 |
| DQA1*05:05:01~DQB1*03:01:01 | 0.0789 | 0.1014 | 0.0831 | 0.0730 | 0.0244 | 0.0441 |
| B*58:02:01~C*06:02:01 | 0.0787 | 0.0627 | 0.1021 | 0.0897 | 0.0915 | 0.0504 |
| DQA1*01:01:02~DQB1*05:01:01 | 0.0770 | 0.0867 | 0.0920 | 0.0730 | 0.0305 | 0.0404 |
| B*53:01:01~C*04:01:01 | 0.0749 | 0.0324 | 0.0828 | 0.1175 | 0.0732 | 0.1437 |
| DQA1*01:05:01~DQB1*05:01:01 | 0.0721 | 0.0583 | 0.0831 | 0.0955 | 0.0610 | 0.0772 |
| DRB1*03:01:01~DRB3*02:02:01 | 0.0700 | 0.1043 | 0.0402 | 0.0506 | 0.0622 | 0.0699 |
| DPB1*105:01:01~DQB1*06:02:01 | 0.0688 | 0.0688 | 0.0698 | 0.0694 | 0.1138 | 0.0456 |
| DRB1*03:02:01~DRB3*01:62:01 | 0.0673 | 0.0610 | 0.0714 | 0.0730 | 0.0938 | 0.0551 |
| DQA1*05:01:01~DQB1*02:01:01 | 0.0666 | 0.0962 | 0.0401 | 0.0449 | 0.0671 | 0.0662 |
| DQB1*02:01:01~DRB1*03:01:01 | 0.0661 | 0.0962 | 0.0401 | 0.0449 | 0.0671 | 0.0625 |
| DPA1*02:01:08~DPB1*01:01:01 | 0.0661 | 0.0374 | 0.0801 | 0.0674 | 0.0854 | 0.0971 |
| DQA1*04:01:01~DQB1*04:02:01 | 0.0627 | 0.0555 | 0.0712 | 0.0674 | 0.0854 | 0.0441 |
| DQB1*05:01:01~DRB1*01:02:01 | 0.0617 | 0.0705 | 0.0638 | 0.0730 | 0.0305 | 0.0441 |
| DQB1*04:02:01~DRB1*03:02:01 | 0.0617 | 0.0556 | 0.0682 | 0.0674 | 0.0854 | 0.0441 |
| DQB1*06:02:01~DRB1*11:01:02 | 0.0614 | 0.0470 | 0.0644 | 0.0899 | 0.0854 | 0.0641 |
| DPA1*01:03:01~DPB1*18:01:01 | 0.0612 | 0.0281 | 0.0668 | 0.1404 | 0.0549 | 0.0863 |
| DRB1*01:02:01~DRB3/4/5*00:00:00 | 0.0609 | 0.0678 | 0.0640 | 0.0730 | 0.0313 | 0.0441 |
| B*42:01:01~C*17:01:01 | 0.0607 | 0.0602 | 0.0666 | 0.0562 | 0.0488 | 0.0576 |
| DPB1*01:01:01~DRB1*03:02:01 | 0.0585 | 0.0622 | 0.0557 | 0.0463 | 0.0842 | 0.0425 |
| DPA1*01:03:01~DPB1*04:01:01 | 0.0582 | 0.0738 | 0.0549 | 0.0337 | 0.0671 | 0.0360 |
| B*15:03:01~C*02:10:01 | 0.0582 | 0.0642 | 0.0725 | 0.0337 | 0.0427 | 0.0324 |
| DPB1*01:01:01~DQB1*04:02:01 | 0.0508 | 0.0479 | 0.0541 | 0.0294 | 0.0793 | 0.0392 |
| Haplotype pairs in the overall African population with frequencies >5% |  |  |  |  |  |  |

We also report haplotypes of class I loci paired with DRB1 frequent in East and West African populations. Notably, haplotype C*04:01:01∼DRB1*11:01:02 was frequent in Tanzania (8.3%) and Uganda (5.9%), while haplotypes A*36:01:01∼DRB1*11:01:02 (7.9%), B*53:01:01∼DRB1*11:01:02 (6.6%), C*06:02:01∼DRB1*15:03:01 (5.4%), and A*02:01:01∼DRB1*15:03:01 (5.1%) were frequent in Tanzania (Supplementary Table 5). In West African Nigeria, C*04:01:01∼DRB1*15:03:01 (5.6%) was the only population-level frequent class I and DRB1 combined haplotype found (Supplementary Table 5).

Additionally, we report 7 multi-locus haplotypes with population-specific frequencies >5% (Supplementary Table 5). Class I haplotypes included A*36:01:01∼B*53:01:01∼C*04:01:01 (8.4%) and A*68:02:01∼B*15:10:01∼C*03:04:02 (5.0%) in Tanzania, and A*02:05:01∼B*58:01:01∼C*07:18:01 (5.5%) in Mozambique (Supplementary Table 5). Notably, class II haplotypes included DPA1*02:02:02∼DPB1*01:01:01∼DQA1*04:01:01∼DQB1*04:02:01∼DRB1*03:02:01∼DRB3* 01:62:01 (8.1%) in Mozambique, while DPA1*01:03:01∼DPB1*02:01:02∼DQA1*01:02:01∼DQB1*06:02:01∼DRB1*15:03:01∼DRB5* 01:01:01 was found in Nigeria (6.8%), Tanzania (6.0%) and Uganda (5.4%) (Supplementary Table 5). In Tanzania, the class II multi-locus population-specific haplotype DPA1*02:02:02∼DPB1*01:01:01∼DQA1*05:05:01∼DQB1*03:19:01∼DRB1*11:01:02∼DRB3* 02:02:01 (5.1%) was frequent (Supplementary Table 5). In Mozambique, the class II multi-locus population-specific haplotype DPA1*03:01:01∼DPB1*105:01:01∼DQA1*01:02:01∼DQB1*06:02:01∼DRB1*15:03:01∼DRB5 *01:01:01 (6.9%) was frequent (Supplementary Table 5). No haplotype with all loci were found at >5% frequency in any population.

### 3.3 Rare, Unique, and Novel alleles in African populations

Alleles with frequencies of <1% were considered rare. HLA class I rare alleles in the overall African population included 26 HLA-A (7.3%), 51 HLA-B (13.9%), and 27 HLA-C (6.5%) alleles (Tables 1-3). HLA class II rare alleles in the African population included 23 DPA1 (5.3%), 50 DPB1 (8.4%), 13 DQA1 (3.3%), 15 DQB1 (3.6%), 25 DRB1 (3.5%), 3 DRB3 (0.2%), 2 DRB4 (0.1%), and 3 DRB5 alleles (0.3%) (Tables 4-6, Supplementary Tables 1-3). Unique population-specific alleles were observed in Kenya (46), Nigeria (36), Uganda (31), Mozambique (13), and Tanzania (6) (Supplementary Table 6). Moreover, East African countries share a specific set of 18 unique alleles, most notably including 11 class I loci A*01:03:01, A*01:09:01, A*02:14, A*29:01:01, A*74:03:01, B*35:02:01, B*44:15:01, B*48:05:01, B*50:01:01, B*73:01:01, and C*04:07:01 (Tables 1-3). In West African Nigeria, the greatest number of unique alleles was observed within HLA-B (8), DPB1 (7), and DRB1 (6) loci, with most unique Nigerian population alleles found in class II loci (21) (Supplementary Table 6). Overall, Kenya had the highest occurrences of unique alleles including B*57:01:01 (9), DPA1*01:30 (9), and DPB1*02:01:22 (9), followed by Nigeria with B*52:01:02 (6) (Supplementary Table 6).

A total of 40 potentially novel alleles were identified during the course of this study, of which 15 novel allele sequences were validated by others in the IPD-IMGT/HLA database prior to this submission (Table 8, Supplementary Table 7). We also extended sequences of one previously published allele where phasing was possible (Supplementary Table 7). We report 25 novel alleles with 19 non-synonymous and 6 synonymous substitutions in exons for HLA-A, B, C, DPA1, DPB1, DQA1, DRB1, and DRB3 (Table 8). Four of the novel alleles had non-synonymous changes in exon 2 which encodes the peptide binding region of the HLA molecule, for loci B, C, DQA1, and DPB1, further supporting that HLA genotyping in these populations is warranted.

**Table 8.** New allele occurrences in the overall Africa cohort.

| Locus | Closest allele | Nucleotide Change | Amino Acid Change | Exon | Occurrence | Population | GenBank Accession |
| --- | --- | --- | --- | --- | --- | --- | --- |
| A | 02:02:01 | 14 C>T | -20 A>V | 1 | 1 | Kenya | PV927387 |
| A | 30:02:01 | 28 C>G | -15 L>V | 1 | 1 | Nigeria | PX239926 |
| B | 15:610 | 47 T>C | -9 V>A | 1 | 1 | Uganda | PX239927 |
| B | 58:01:01 | 175 A>G | 35 R>G | 2 | 1 | Kenya | PX239928 |
| C | 07:01:01 | 197 G>A | 42 S>N | 2 | 1 | Nigeria | PX239929 |
| DPA1 | 02:01:01 | 153 G>A | 20 G | 2 | 1 | Nigeria | PX239930 |
| DPA1 | 02:09:01 | 763 C>T | 224 R>W | 4 | 1 | Kenya | PX239931 |
| DPA1 | 03:01:01 | 49 G>A | -15 A>T | 1 | 1 | Uganda | PX239933 |
| DPA1 | 03:01:01 | 437 C>T | 115 P>L | 3 | 2 | Kenya, Uganda | PX239932 |
| DPB1 | 02:01:02 | 588 T>C | 167 D | 3 | 1 | Uganda | PX239934 |
| DPB1 | 02:01:19 | 81 C>T | -3 V | 1 | 1 | Mozambique | PX239935 |
| DPB1 | 40:01:01 | 667 C>T | 194 R>W | 4 | 1 | Uganda | PX239937 |
| DPB1 | 49:01:01 | 280 A>C | 65 I>L | 2 | 1 | Uganda | PX239938 |
| DPB1 | 75:01 | 619 A>C | 178 M>L | 3 | 2 | Kenya, Uganda | PX239939 |
| DPB1 | 135:01:01 | 668 G>A | 194 R>Q | 4 | 1 | Kenya | PX239936 |
| DQA1 | 01:02:01 | 295 A>G | 76 M>V | 2 | 1 | Nigeria | PX239941 |
| DQA1 | 01:02:01 | 496 A>G | 143 K>E | 3 | 3 | Kenya | PX239940 |
| DQA1 | 01:05:01 | 333 G>A | 88 E | 3 | 2 | Kenya | PX239942 |
| DRB1 | 01:01:01 | 705 C>T | 206 F | 4 | 1 | Uganda | PX239943 |
| DRB1 | 03:01:01 | 620 C>A | 178 P>Q | 3 | 1 | Kenya | PX239944 |
| DRB1 | 04:10:03 | 609 A>C | 174 Q>H | 3 | 1 | Kenya | PX239945 |
| DRB1 | 14:54:01 | 653 G>A | 189 R>K | 4 | 2 | Mozambique, Uganda | PX239947 |
| DRB1 | 15:03:01 | 725 T>C | 213 L>P | 4 | 1 | Tanzania | PX239948 |
| DRB1 | 11:01:02 | 733 G>A | 216 G>R | 4 | 1 | Uganda | PX239946 |
| DRB3 | 02:02:01 | 393 T>C | 102 Y | 3 | 2 | Tanzania | PX239949 |

Novel alleles identified in the African cohorts. The closest HLA reference sequences for each allele are shown for synonymous and non-synonymous exonic variants. Occurrence in cohort represents the number of times this allele was found in this dataset.

## Discussion

We conducted large-scale NGS HLA genotyping in African cohorts for deeper characterization of HLA genotype profiles from greater than 1000 individuals from Kenya, Uganda, Tanzania, Mozambique, and Nigeria. Advances in NGS technologies have enabled us to achieve 6-fold greater sequencing depth per locus while increasing throughput of multiplexed donors to 384 in a single sequencing run. This, combined with our previously published MiSeq NGS-HLA genotyping approach, provided higher resolution which contributed to the discovery of 25 novel alleles and revealed several unique and rare population HLA alleles^34-37,39^. We previously identified DPB1*665:01 in one participant of unknown ancestry, and now find this allele in the Kenyan and Ugandan populations^37^. Overall, allelic diversity aligned with increased sample size observed in ascending order of each population and HLA-B loci had the greatest number of alleles regardless of geographical location.

There are very few published NGS-based HLA genotyping data from Africa, and our study is the first effort to characterize all 11 HLA loci using contiguous full-length gene sequencing. Given the limited availability of high resolution NGS-based HLA genotyping in African populations, we compared genotype frequencies in our dataset at 2-field resolution to match with other reported NGS and medium-resolution HLA data (Polymerase Chain Reaction with Sequence-Specific Oligonucleotide Probes (PCR-SSOP)/Sanger Sequencing-Based Typing (SBT)) (Supplementary Table 8). We reviewed these studies and datasets for total unique, rare, and frequent population alleles at 2-field resolution in our cohorts and it followed the same sample size-dependent relationship as observed at 3-field. When compared to other African studies of various sample sizes we observed both similarities and unique or rare differences across population HLA diversity. Collectively, this allowed us to assess the impact of deeper, expanded genotyping in African countries.

The Kenyan cohort with a sample size of 375 was the largest of the populations in this study, and subsequently the most genetically diverse at all class I and DP, DQB1and DRB3 loci. Arlehamn et al. (n=100) and Banjoko et al. (n=109) conducted NGS genotyping of various loci in the Kenyan population and identified fewer alleles, as expected from studies of smaller sample size^37,41^. In our study we identified HLA-B*49:01 and C*07:18 as common alleles at frequencies >5%, which has not been the case previously. Together, these results highlight the increased allelic resolution and diversity achievable through larger, high resolution NGS studies. The second largest cohort, from Uganda (n=338), had a total of 115 and 116 class I and II alleles, respectively, at the 2-field level and showed the greatest diversity at the DQA1 locus when compared to the other countries in this study. Digital et al. reported exon-based SBT typing of 8 HLA loci in 892 Ugandans, reporting a total of 124 class I alleles and 102 class II alleles (no DRB3/4/5 which are not reported)^53^. They observed a greater number of HLA alleles, as expected from their much larger sample size. Further, we observe DPB1*105:01 at frequencies of >10% in Uganda which was not reported in Digital et al.^53^, likely due to genotyping restricted to specific exons potentially leading to ambiguities. However, a higher frequency of DPB1*105:01 has also been observed in South Africa^54^, highlighting the need to expand NGS based HLA genotyping in Ugandan populations.

The third largest population was the West African Nigerian cohort (n=139) with 91 class I and 94 class II 2-field alleles. This cohort was the most diverse at the DRB1 locus when compared to the other African cohorts. Testi et al. previously genotyped 97 Nigerians by SSOP/SBT, identifying 25 HLA-A, 31 B, 22 C, 16 DQB1, and 22 DRB1 alleles at 2-field resolution^55^. In comparison, our study reports slightly higher allelic diversity for all loci except DQB1 (30 A, 35 B, 26 C, 16 DQB1, 24 DRB1) at 2-field resolution, and frequencies were largely consistent between our studies^55^. However, this is the first report of NGS-based HLA genotyping in a Nigerian population, and is also the first genotyping data for DQA1 and DRB3/4/5 loci^32^. In accordance with the representative allelic diversity that a larger cohort may have captured, expanded genotyping in the Nigerian population may reveal a greater diversity at other loci besides DRB1.

The cohorts from Tanzania (n=89) and Mozambique (n=82) were smaller than those from the other three African countries. Nevertheless, this is the first comprehensive report of alleles at all HLA class II loci in Tanzania. Barton et al. captured a larger number of HLA alleles using NGS in 336 Tanzanian individuals but for only a subset of HLA loci^56^. Despite our smaller cohort of 89 individuals, we found 22 alleles not reported previously, including one allele with a frequency >5% (DPB1*104:01), three alleles with frequencies between 1% and 5%, and 17 rare alleles across class I and class II loci^56^. These results highlight unique allelic diversity and emphasize the need for broader NGS-based HLA genotyping in Tanzania. In the other smaller cohort from Mozambique, we identified 74 class I and 86 class II alleles at the 2-field level. To our knowledge, our study is not only the first to employ NGS genotyping for HLA-C, DPA1, DPB1, DQA1, DQB1, and DRB3/4/5 in Mozambique, but also the first to generate >1-field resolution typing, unlike previously published cohorts^32,57^. More HLA typing of participants from Mozambique using NGS will be necessary for deeper characterization.

In addition to characterizing HLA alleles in a population, the frequencies of previously reported alleles shown to have associations with HIV viral load control and disease progression were assessed. HLA alleles known to be protective against HIV, and validated in multiple cohorts, include B*13:02, B*14:02, B*27:05, B*44:03, B*57:01, B*57:03, B*58:01, B*81:01, A*25:01, A*32:01, and A*74:01^29^. Other than A*25:01, all these alleles were detected in our cohorts. HLA-B*57 is known to have the strongest association with viral control in different world populations^58-60^. B*57:03 is the prevalent B*57 subtype in African populations, and has a known protective role against HIV disease progression^61-63^. This African-specific subtype was present in participants from all the African countries typed in this study, but at frequencies of <5%. As expected, we did not detect B*57:03 in the Philippine and Thai populations that we assessed previously with the same approach^35,39^. Interestingly, the B*57:01 subtype common to countries of European ancestry was observed with a frequency of 1.2% in Kenya, which was between the 0.57% and 1.35% observed in the Philippine and Thai populations, respectively, confirming the low prevalence of B*57:01 outside of populations with European ancestry^58,59^. Another notable protective allele, B*58:01, was detected at a remarkably high frequency in Mozambique (10.4%) and present in all the other African countries with frequencies >5%. B*81:01, another African-specific allele, was found in all the African countries studied here with an overall presence of 3.8%, but was absent in the two Southeast Asian countries^35,39,64^. B*27:05 is associated with slower HIV progression, and was found in a single individual in these African cohorts, mirroring its rarity in the Southeast Asian countries^22,23,35,39,65,66^. HLA alleles that associate with poor HIV disease outcomes, including B*08:01, B*18:01, B*35-Px, B*58:02, B*45:01, and A*36:01, were all observed in our African cohorts^24,25,27,60,67-69^. B*18:01 was found at frequencies of 6.7%, 3.3%, 3%, and 1.5% in Mozambique, total African, Thai, and Philippine cohorts, respectively^27,35,39^. B*53:01, a subtype of B*35-Px previously shown to associate with increased disease progression and high viral loads, was found at high frequencies in all these countries; notably this allele is in the B*53:01:01∼C*04:01:01 haplotype which is the most frequent class I haplotype in Tanzania and Nigeria populations^24^. Similarly, B*58:02, an allele that associates with higher viral load and faster disease progression, was observed with higher frequencies in all the African countries, with B*58:02:01∼C*06:02:01 being the most common class I haplotype^35,39,69,70^. B*45:01, an allele which strongly associates with higher viral load in central and south African populations, was found at the highest frequencies within the Tanzania cohort (11.2%)^18,35,39,60,67^. However, alleles known to associate with HIV outcomes in other world populations may not reflect associations in Africa because of differences in HLA types across populations and viral adaptations. We previously reported a novel association of B*46:01 with increased disease progression in Thailand in three independent cohorts^28^. Since B*46:01 is only prevalent in Southeast Asia, this discovery, more than 25 years after the first report of HLA genetic associations, speaks to the importance of large-scale NGS characterization of HLA in countries outside the Western hemisphere.

Our haplotype analyses also support a high level of diversity in African countries with very few common haplotypes for HLA class I. Notably, an overrepresentation of a class II haplotype DQA1*01:02:01∼DQB1*06:02:01 was observed in all African populations with frequencies ranging from 16.2-29.4%. Previous findings in an HIV-1 efficacy study in Thailand showed that the DQB1*06 allele associates with reduced vaccine protection^30^. Interestingly in the same study, the DPB1*13:01 allele associated with increased HIV-1 specific antibody titers and higher vaccine efficacy, but its absence in African populations may have contributed to the ineffectiveness of the same vaccine in the continent^71,72^. Despite the paucity of HIV clinical trials with efficacy data, HLA class II alleles have been shown to associate with vaccine protection and antibody titers in other trials in Africa^73,74^. We also describe other class II alleles such as DPB1*01:01, DPB1*105:01, DRB1*15:03, DRB3*02:02, and DRB5*01:01 as very frequent alleles (>20%) in African populations which were not previously reported across other similar and global populations ^20,53,55,56,75^.

Irrespective of Africa’s extraordinary genetic diversity, high-resolution HLA genotyping studies in these populations, using next-generation sequencing of all HLA loci, have been limited to date^31,32^. Our study contributes to narrowing this gap by reporting one of the most comprehensive high-resolution, NGS-based HLA genotyping efforts in African populations, characterizing allele and haplotype frequencies across 11 loci from over 1000 individuals in East, Southeast, and West African countries. We report the first full-length class II genotyping data from participants in Kenya, Uganda, and Tanzania, and the first high-resolution HLA data from participants in Mozambique and Nigeria^32^. Our findings reveal extensive allelic diversity and pronounced intra-continental variation, particularly at the highly polymorphic HLA-B locus, as well as novel allelic diversity across both class I and II loci. For three of the novel alleles, although the mismatched position could not be fully phased with the other exons, the single-nucleotide variation for these alleles were mostly phased due to sequencing of introns, which supported the final determination of the novel allele status and emphasizes the importance of intronic sequences in HLA genotyping.

Further, we observed that overall the DQB1 locus was not in HWE and that the expected DQA1∼DQB1 haplotypes were not present for all participants when homozygous for DQB1*05, DQB1*06, or DQB1*02. Allele dropout due to novel alleles at primer binding sites could be the main cause, as the loss in heterozygosity was mainly detected in Kenya. The presence of differential DNA methylation, G-quadruplex structures, homopolymeric tracts, and pseudogenes could be other factors that warrant further investigation in future studies^76,77^.

The diversity found within HLA class I loci in the African cohort is functionally significant, as individuals heterozygous at class I loci are known to have a broader range of T-cell recognition, which is associated with better control of HIV viral replication and slower disease progression ^11,14,25,64^. Together, the combined insights from both class I and class II genotyping highlight the immunogenic complexity of African populations and the need for comprehensive, population-specific reference datasets. Our study provides essential data for improving population-informed epitope prediction algorithms, enhancing HIV vaccine designs, and advancing our understanding of host-pathogen interactions in infectious disease.

## Acknowledgments

This work was supported by agreements #W81XWH-18-2-0040 and #HT94252430004 between the Henry M. Jackson Foundation for the Advancement of Military Medicine, Inc., and the U.S. Department of Defense (DOD). This work was supported by cooperative agreements #W81XWH-18-2-0040 and # HT94252420020 between the Henry M. Jackson Foundation for the Advancement of Military Medicine, Inc., and the U.S. Department of Defense (DOD). This research has been supported by the President’s Emergency Plan for AIDS Relief (PEPFAR) through the U.S. DOD. This research was funded, in part, by the U.S. National Institute of Allergy and Infectious Diseases. The views expressed are those of the authors and should not be construed to represent the positions of the U.S. Army or the U.S. Department of Defense (DOD). The investigators have adhered to the policies for protection of human participants as prescribed in AR 70–25.

We thank the RV217, RV251, RV329, RV456 and RV460 study volunteers and study teams in Kenya, Uganda, Tanzania, Mozambique, and Nigeria for their contributions and commitment to HIV and immunological clinical research.

## Data Availability Statement

The data that supports the findings of this study are available on request from corresponding author.

## Author contributions

RT conceptualized and led the study. HK, FW-M, LM, ATi, JM, JK, FS, GRM, MLR, and JAA provided clinical samples and oversight of the studies. YVR, PKE, and ATy performed HLA genotyping. YVR, AG, ATy and RT performed HLA data analysis. YVR, LRI and RT drafted the original manuscript. AG and PKE reviewed and edited the original manuscript. All authors read and reviewed the manuscript.

## Ethics statement

HLA genotyping was approved for all sites by the participating local institutional review boards and in the USA. The investigators have adhered to the policies for protection of human participants as prescribed in AR 70–25.

## References

1. MHC.Sequencing.Consortium. Complete sequence and gene map of a human major histocompatibility complex. Nature. 1999/10/01 1999;401(6756):921-923. doi:10.1038/44853

2. Shiina T, Hosomichi K, Inoko H, Kulski JK. The HLA genomic loci map: expression, interaction, diversity and disease. Journal of Human Genetics. 2009/01/01 2009;54(1):15-39. doi:10.1038/jhg.2008.5

3. Kaslow RA, McNicholl JM. Genetic determinants of HIV-1 infection and its manifestations. Proc Assoc Am Physicians. Jul-Aug 1999;111(4):299–307. doi:10.1046/j.1525-1381.1999.99238.x

4. Migueles SA, Laborico AC, Imamichi H, et al. The differential ability of HLA B*5701+ long-term nonprogressors and progressors to restrict human immunodeficiency virus replication is not caused by loss of recognition of autologous viral gag sequences. J Virol. Jun 2003;77(12):6889–98. doi:10.1128/jvi.77.12.6889-6898.2003

5. Fellay J, Shianna KV, Ge D, et al. A whole-genome association study of major determinants for host control of HIV-1. Science. Aug 17 2007;317(5840):944-7. doi:10.1126/science.1143767

6. Thomas R, Apps R, Qi Y, et al. HLA-C cell surface expression and control of HIV/AIDS correlate with a variant upstream of HLA-C. Nat Genet. Dec 2009;41(12):1290–4. doi:10.1038/ng.486

7. Gingras SN, Tang D, Tuff J, McLaren PJ. Minding the gap in HIV host genetics: opportunities and challenges. Hum Genet. Jun 2020;139(6-7):865–875. doi:10.1007/s00439-020-02177-9

8. McLaren PJ, Carrington M. The impact of host genetic variation on infection with HIV-1. Nat Immunol. Jun 2015;16(6):577–83. doi:10.1038/ni.3147

9. Culmann B, Gomard E, Kiény MP, et al. Six epitopes reacting with human cytotoxic CD8+ T cells in the central region of the HIV-1 NEF protein. J Immunol. Mar 1 1991;146(5):1560–5.

10. Koshino K, Tokano Y, Hishikawa T, Sekigawa I, Takasaki Y, Hashimoto H. Detection of antibodies to HIV-1 gp41- and HLA class II antigen-derived peptides in SLE patients. Scand J Rheumatol. 1995;24(5):288–92. doi:10.3109/03009749509095165

11. Carrington M, Nelson GW, Martin MP, et al. HLA and HIV-1: heterozygote advantage and B*35-Cw*04 disadvantage. Science. Mar 12 1999;283(5408):1748-52. doi:10.1126/science.283.5408.1748

12. Viard M, O’HUigin C, Yuki Y, et al. Impact of HLA class I functional divergence on HIV control. Science. Jan 19 2024;383(6680):319-325. doi:10.1126/science.adk0777

13. World Health Orgnaization Africa Region, Health topics, HIV/AIDS.

14. Rousseau CM, Daniels MG, Carlson JM, et al. HLA class I-driven evolution of human immunodeficiency virus type 1 subtype c proteome: immune escape and viral load. J Virol. Jul 2008;82(13):6434–46. doi:10.1128/jvi.02455-07

15. Patel PH, Preston BD. Marked infidelity of human immunodeficiency virus type 1 reverse transcriptase at RNA and DNA template ends. Proc Natl Acad Sci U S A. Jan 18 1994;91(2):549–53. doi:10.1073/pnas.91.2.549

16. Moore CB, John M, James IR, Christiansen FT, Witt CS, Mallal SA. Evidence of HIV-1 adaptation to HLA-restricted immune responses at a population level. Science. May 24 2002;296(5572):1439-43. doi:10.1126/science.1069660

17. Kawashima Y, Pfafferott K, Frater J, et al. Adaptation of HIV-1 to human leukocyte antigen class I. Nature. Apr 2 2009;458(7238):641-5. doi:10.1038/nature07746

18. Cao K, Moormann AM, Lyke KE, et al. Differentiation between African populations is evidenced by the diversity of alleles and haplotypes of HLA class I loci. Tissue Antigens. 2004;63(4):293–325. doi:10.1111/j.0001-2815.2004.00192.x

19. Gomez F, Hirbo J, Tishkoff SA. Genetic variation and adaptation in Africa: implications for human evolution and disease. Cold Spring Harb Perspect Biol. Jul 1 2014;6(7):a008524. doi:10.1101/cshperspect.a008524

20. Sanchez-Mazas A, Nunes JM. The most frequent HLA alleles around the world: A fundamental synopsis. Best Pract Res Clin Haematol. Jun 2024;37(2):101559. doi:10.1016/j.beha.2024.101559

21. Farinre O, Gounder K, Reddy T, et al. Subtype-specific differences in Gag-protease replication capacity of HIV-1 isolates from East and West Africa. Retrovirology. May 5 2021;18(1):11. doi:10.1186/s12977-021-00554-4

22. Kaslow RA, Carrington M, Apple R, et al. Influence of combinations of human major histocompatibility complex genes on the course of HIV-1 infection. Nat Med. Apr 1996;2(4):405–11. doi:10.1038/nm0496-405

23. Fellay J, Ge D, Shianna KV, et al. Common genetic variation and the control of HIV-1 in humans. PLoS Genet. Dec 2009;5(12):e1000791. doi:10.1371/journal.pgen.1000791

24. Gao X, Nelson GW, Karacki P, et al. Effect of a single amino acid change in MHC class I molecules on the rate of progression to AIDS. N Engl J Med. May 31 2001;344(22):1668–75. doi:10.1056/nejm200105313442203

25. Kiepiela P, Leslie AJ, Honeyborne I, et al. Dominant influence of HLA-B in mediating the potential co-evolution of HIV and HLA. Nature. Dec 9 2004;432(7018):769-75. doi:10.1038/nature03113

26. Apps R, Qi Y, Carlson JM, et al. Influence of HLA-C expression level on HIV control. Science. Apr 5 2013;340(6128):87-91. doi:10.1126/science.1232685

27. Leslie A, Matthews PC, Listgarten J, et al. Additive contribution of HLA class I alleles in the immune control of HIV-1 infection. J Virol. Oct 2010;84(19):9879–88. doi:10.1128/jvi.00320-10

28. Li SS, Hickey A, Shangguan S, et al. HLA-B∗46 associates with rapid HIV disease progression in Asian cohorts and prominent differences in NK cell phenotype. Cell Host & Microbe. 2022/08/10/ 2022;30(8):1173-1185.e8. 10.1016/j.chom.2022.06.005

29. Goulder PJ, Walker BD. HIV and HLA class I: an evolving relationship. Immunity. Sep 21 2012;37(3):426–40. doi:10.1016/j.immuni.2012.09.005

30. Prentice HA, Tomaras GD, Geraghty DE, et al. HLA class II genes modulate vaccine-induced antibody responses to affect HIV-1 acquisition. Science translational medicine. Jul 15 2015;7(296):296ra112. doi:10.1126/scitranslmed.aab4005

31. Arrieta-Bolaños E, Hernández-Zaragoza DI, Barquera R. An HLA map of the world: A comparison of HLA frequencies in 200 worldwide populations reveals diverse patterns for class I and class II. Front Genet. 2023;14:866407. doi:10.3389/fgene.2023.866407

32. Gonzalez-Galarza FF, McCabe A, Santos E, et al. Allele frequency net database (AFND) 2020 update: gold-standard data classification, open access genotype data and new query tools. Nucleic Acids Res. Jan 8 2020;48(D1):D783–d788. doi:10.1093/nar/gkz1029

33. Prentice HA, Ehrenberg PK, Baldwin KM, et al. HLA class I, KIR, and genome-wide SNP diversity in the RV144 Thai phase 3 HIV vaccine clinical trial. Immunogenetics. 2014/05/01 2014;66(5):299-310. doi:10.1007/s00251-014-0765-6

34. Ehrenberg PK, Geretz A, Thomas R. High-Throughput Contiguous Full-Length Next-Generation Sequencing of HLA Class I and II Genes from 96 Donors in a Single MiSeq Run. Methods Mol Biol. 2018;1802:89–100. doi:10.1007/978-1-4939-8546-3_6

35. Geretz A, Cofer L, Ehrenberg PK, et al. Next-generation sequencing of 11 HLA loci in a large dengue vaccine cohort from the Philippines. Hum Immunol. Aug 2020;81(8):437–444. doi:10.1016/j.humimm.2020.06.010

36. Ehrenberg PK, Geretz A, Baldwin KM, et al. High-throughput multiplex HLA genotyping by next-generation sequencing using multi-locus individual tagging. BMC Genomics. Oct 06 2014;15:864. doi:10.1186/1471-2164-15-864

37. Ehrenberg PK, Geretz A, Sindhu RK, et al. High-throughput next-generation sequencing to genotype six classical HLA loci from 96 donors in a single MiSeq run. HLA. Nov 2017;90(5):284–291. doi:10.1111/tan.13133

38. Baldwin KM, Ehrenberg PK, Geretz A, et al. HLA class II diversity in HIV-1 uninfected individuals from the placebo arm of the RV144 Thai vaccine efficacy trial. Tissue Antigens. Feb 2015;85(2):117–26. doi:10.1111/tan.12507

39. Geretz A, Ehrenberg PK, Bouckenooghe A, et al. Full-length next-generation sequencing of HLA class I and II genes in a cohort from Thailand. Hum Immunol. Nov 2018;79(11):773–780. doi:10.1016/j.humimm.2018.09.005

40. Robb ML, Eller LA, Kibuuka H, et al. Prospective Study of Acute HIV-1 Infection in Adults in East Africa and Thailand. N Engl J Med. Jun 2 2016;374(22):2120–30. doi:10.1056/NEJMoa1508952

41. Graham BS, Koup RA, Roederer M, et al. Phase 1 safety and immunogenicity evaluation of a multiclade HIV-1 DNA candidate vaccine. J Infect Dis. Dec 15 2006;194(12):1650–60. doi:10.1086/509259

42. Ake JA, Polyak CS, Crowell TA, et al. Noninfectious Comorbidity in the African Cohort Study. Clin Infect Dis. Aug 1 2019;69(4):639–647. doi:10.1093/cid/ciy981

43. Ake JA, Paolino K, Hutter JN, et al. Safety and Immunogenicity of an Accelerated Ebola Vaccination Schedule in People with and without Human Immunodeficiency Virus: A Randomized Clinical Trial. Vaccines (Basel). May 4 2024;12(5)doi:10.3390/vaccines12050497

44. Mwesigwa B, Sawe F, Oyieko J, et al. Safety and Immunogenicity of Accelerated Heterologous Two-dose Ebola Vaccine Regimens in Adults With and Without HIV in Africa. Clin Infect Dis. Apr 24 2024;doi:10.1093/cid/ciae215

45. Alving CR, K. PK, R. MG, Mangala R, and Beck Z. Army Liposome Formulation (ALF) family of vaccine adjuvants. Expert Review of Vaccines. 2020/03/03 2020;19(3):279-292. doi:10.1080/14760584.2020.1745636

46. Osoegawa K, Vayntrub TA, Wenda S, et al. Quality control project of NGS HLA genotyping for the 17th International HLA and Immunogenetics Workshop. Hum Immunol. Apr 2019;80(4):228–236. doi:10.1016/j.humimm.2019.01.009

47. Robinson J, Barker DJ, Marsh SGE. 25 years of the IPD-IMGT/HLA Database. HLA. 2024;103(6):e15549. 10.1111/tan.15549

48. Barker DJ, Maccari G, Georgiou X, et al. The IPD-IMGT/HLA Database. Nucleic Acids Research. 2022;51(D1):D1053-D1060. doi:10.1093/nar/gkac1011

49. Vegan: Community Ecology Package. R package version 2.5-7. 2020.

50. Guo SW, Thompson EA. Performing the exact test of Hardy-Weinberg proportion for multiple alleles. Biometrics. Jun 1992;48(2):361–72.

51. Watterson GA. The homozygosity test of neutrality. Genetics. Feb 1978;88(2):405–17.

52. Guess HA, Ewens WJ. Theoretical and simulation results relating to the neutral allele theory. Theor Popul Biol. Dec 1972;3(4):434–47. doi:10.1016/0040-5809(72)90015-9

53. Digitale JC, Callaway PC, Martin M, et al. HLA Alleles B(*)53:01 and C(*)06:02 Are Associated With Higher Risk of P. falciparum Parasitemia in a Cohort in Uganda. Front Immunol. 2021;12:650028. doi:10.3389/fimmu.2021.650028

54. Thorstenson YR, Creary LE, Huang H, et al. Allelic resolution NGS HLA typing of Class I and Class II loci and haplotypes in Cape Town, South Africa. Hum Immunol. Dec 2018;79(12):839–847. doi:10.1016/j.humimm.2018.09.004

55. Testi M, Battarra M, Lucarelli G, et al. HLA-A-B-C-DRB1-DQB1 phased haplotypes in 124 Nigerian families indicate extreme HLA diversity and low linkage disequilibrium in Central-West Africa. Article. Tissue Antigens. 2015;86(4):285–292. doi:10.1111/tan.12642

56. Barton A, Ramadhani A, Mafuru E, et al. HLA-A, -B, -C, -DPB1, -DQB1 and -DRB1 allele frequencies of North Tanzanian Maasai. Hum Immunol. Feb 2023;84(2):67-68. doi:10.1016/j.humimm.2022.10.008

57. Assane AAA, Fabricio-Silva GM, Cardoso-Oliveira J, et al. Human leukocyte antigen–A, –B, and –DRB1 allele and haplotype frequencies in the Mozambican population: A blood donor–based population study. Human Immunology. 2010/10/01/ 2010;71(10):1027-1032. 10.1016/j.humimm.2010.06.017

58. Kolou M, Poda A, Diallo Z, et al. Prevalence of human leukocyte antigen HLA-B*57:01 in individuals with HIV in West and Central Africa. BMC Immunology. 2021/07/22 2021;22(1):48. doi:10.1186/s12865-021-00427-7

59. Martin MA, Klein TE, Dong BJ, Pirmohamed M, Haas DW, Kroetz DL. Clinical pharmacogenetics implementation consortium guidelines for HLA-B genotype and abacavir dosing. Clin Pharmacol Ther. Apr 2012;91(4):734–8. doi:10.1038/clpt.2011.355

60. Tang J, Tang S, Lobashevsky E, et al. Favorable and unfavorable HLA class I alleles and haplotypes in Zambians predominantly infected with clade C human immunodeficiency virus type 1. J Virol. Aug 2002;76(16):8276–84. doi:10.1128/jvi.76.16.8276-8284.2002

61. McKinnon LR, Capina R, Peters H, et al. Clade-specific evolution mediated by HLA-B*57/5801 in human immunodeficiency virus type 1 clade A1 p24. J Virol. Dec 2009;83(23):12636–42. doi:10.1128/jvi.01236-09

62. Nii-Trebi NI, Matsuoka S, Kawana-Tachikawa A, et al. Super high-resolution single-molecule sequence-based typing of HLA class I alleles in HIV-1 infected individuals in Ghana. PLoS One. 2022;17(6):e0269390. doi:10.1371/journal.pone.0269390

63. Lindquist L, Kilembe W, Karita E, et al. HLA-A*23 Is Associated With Lower Odds of Acute Retroviral Syndrome in Human Immunodeficiency Virus Type 1 Infection: A Multicenter Sub-Saharan African Study. Open Forum Infect Dis. Apr 2024;11(4):ofae129. doi:10.1093/ofid/ofae129

64. Leslie A, Price DA, Mkhize P, et al. Differential Selection Pressure Exerted on HIV by CTL Targeting Identical Epitopes but Restricted by Distinct HLA Alleles from the Same HLA Supertype1. The Journal of Immunology. 2006;177(7):4699–4708. doi:10.4049/jimmunol.177.7.4699

65. McNeil AJ, Yap PL, Gore SM, et al. Association of HLA types A1-B8-DR3 and B27 with rapid and slow progression of HIV disease. Qjm. Mar 1996;89(3):177–85. doi:10.1093/qjmed/89.3.177

66. Schneidewind A, Brockman MA, Yang R, et al. Escape from the dominant HLA-B27-restricted cytotoxic T-lymphocyte response in Gag is associated with a dramatic reduction in human immunodeficiency virus type 1 replication. J Virol. Nov 2007;81(22):12382–93. doi:10.1128/jvi.01543-07

67. Carlson JM, Listgarten J, Pfeifer N, et al. Widespread impact of HLA restriction on immune control and escape pathways of HIV-1. J Virol. May 2012;86(9):5230–43. doi:10.1128/jvi.06728-11

68. Kiepiela P, Ngumbela K, Thobakgale C, et al. CD8+ T-cell responses to different HIV proteins have discordant associations with viral load. Nature Medicine. 2007/01/01 2007;13(1):46-53. doi:10.1038/nm1520

69. Ngumbela KC, Day CL, Mncube Z, et al. Targeting of a CD8 T cell env epitope presented by HLA-B*5802 is associated with markers of HIV disease progression and lack of selection pressure. AIDS Res Hum Retroviruses. Jan 2008;24(1):72–82. doi:10.1089/aid.2007.0124

70. Lazaryan A, Lobashevsky E, Mulenga J, et al. Human leukocyte antigen B58 supertype and human immunodeficiency virus type 1 infection in native Africans. J Virol. Jun 2006;80(12):6056–60. doi:10.1128/jvi.02119-05

71. Zolla-Pazner S, Michael NL, Kim JH. A tale of four studies: HIV vaccine immunogenicity and efficacy in clinical trials. Lancet HIV. Jul 2021;8(7):e449–e452. doi:10.1016/s2352-3018(21)00073-4

72. Moodie Z, Dintwe O, Sawant S, et al. Analysis of the HIV Vaccine Trials Network 702 Phase 2b–3 HIV-1 Vaccine Trial in South Africa Assessing RV144 Antibody and T-Cell Correlates of HIV-1 Acquisition Risk. The Journal of Infectious Diseases. 2022;226(2):246–257. doi:10.1093/infdis/jiac260

73. Thomas R, Chansinghakul D, Limkittikul K, et al. Associations of human leukocyte antigen with neutralizing antibody titers in a tetravalent dengue vaccine phase 2 efficacy trial in Thailand. Human Immunology. 2022/01/01/ 2022;83(1):53-60. 10.1016/j.humimm.2021.09.006

74. Mentzer AJ, Dilthey AT, Pollard M, et al. High-resolution African HLA resource uncovers HLA-DRB1 expression effects underlying vaccine response. Nat Med. May 2024;30(5):1384–1394. doi:10.1038/s41591-024-02944-5

75. Arlehamn CSL, Copin R, Leary S, et al. Sequence-based HLA-A, B, C, DP, DQ, and DR typing of 100 Luo infants from the Boro area of Nyanza Province, Kenya. Human Immunology. 2017/04/01/ 2017;78(4):325-326. 10.1016/j.humimm.2017.03.007

76. Kong D, Lee N, Dela Cruz ID, et al. Concurrent typing of over 4000 samples by long-range PCR amplicon-based NGS and rSSO revealed the need to verify NGS typing for HLA allelic dropouts. Human Immunology. 2021/08/01/ 2021;82(8):581-587. 10.1016/j.humimm.2021.04.008

77. Stevens AJ, Taylor MG, Pearce FG, Kennedy MA. Allelic Dropout During Polymerase Chain Reaction due to G-Quadruplex Structures and DNA Methylation Is Widespread at Imprinted Human Loci. G3 (Bethesda). Mar 10 2017;7(3):1019-1025. doi:10.1534/g3.116.038687

