## Supplementary data for "Full-length next-generation sequencing of 11 HLA loci of more than 1000 individuals from clinical cohorts in East and West Africa"

### Supplementary Figures

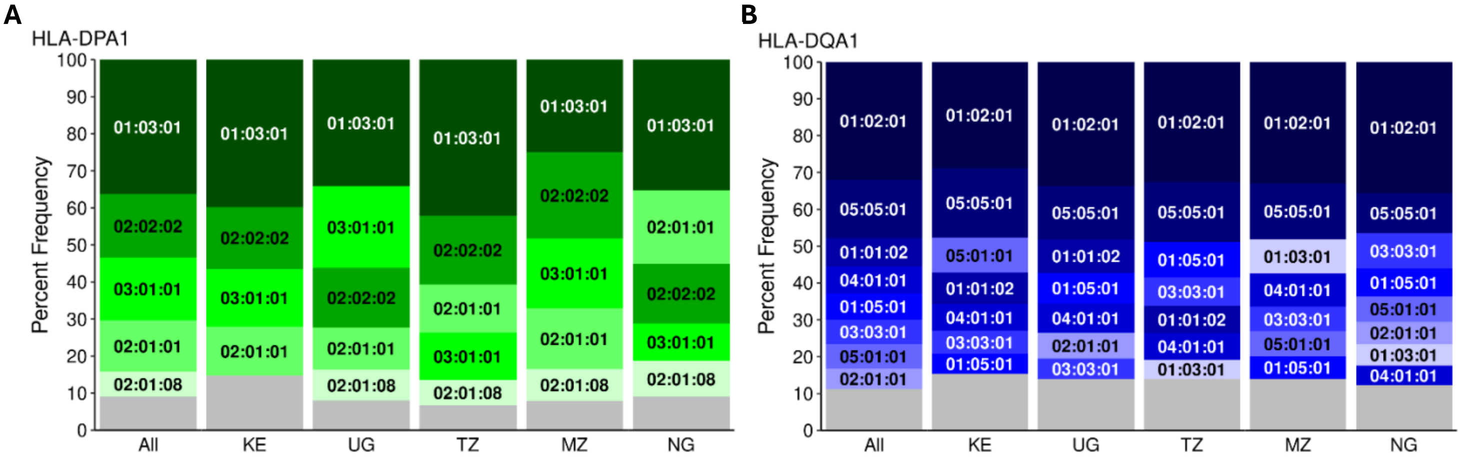

**Supplementary Figure 1.** Common HLA class II frequencies of 1023 individuals in African countries (All), Kenya (KE), Uganda (UG), Tanzania (TZ), Mozambique (MZ), and Nigeria (NG). Frequencies greater than 5% are represented in stacked plots over the sum of remaining low frequency alleles (grey) for HLA class II loci **A.** HLA-DPA1, and **B.** HLA-DQA1

**Supplementary Table 1.** HLA-DPA1 allele frequencies in Africa across East, Southeast, and West countries at 3-field resolution

| | **Africa** | | **East** | | | | | | **Southeast** | | **West** | |
|---|---|---|---|---|---|---|---|---|---|---|---|---|
| | Overall | | Kenya | | Uganda | | Tanzania | | Mozambique | | Nigeria | |
| **Allele** | Frequency | 2N=2046 | Frequency | 2N=750 | Frequency | 2N=676 | Frequency | 2N=178 | Frequency | 2N=164 | Frequency | 2N=278 |
| DPA1*01:03:01 | 0.3632 | 743 | 0.3973 | 298 | 0.3417 | 231 | 0.4214 | 75 | 0.2500 | 41 | 0.3525 | 98 |
| DPA1*01:03:03 | 0.0020 | 4 | 0.0027 | 2 | 0.0015 | 1 | 0.0056 | 1 | | | | |
| DPA1*01:03:10 | 0.0005 | 1 | 0.0013 | 1 | | | | | | | | |
| DPA1*01:04:01 | 0.0083 | 17 | 0.0133 | 10 | 0.0030 | 2 | 0.0056 | 1 | 0.0122 | 2 | 0.0072 | 2 |
| DPA1*01:06:02 | 0.0005 | 1 | 0.0013 | 1 | | | | | | | | |
| DPA1*01:12:01 | 0.0010 | 2 | | | 0.0030 | 2 | | | | | | |
| DPA1*01:19 | 0.0010 | 2 | | | | | | | | | 0.0072 | 2 |
| DPA1*01:30 | 0.0044 | 9 | 0.0120 | 9 | | | | | | | | |
| DPA1*01:58:01 | 0.0005 | 1 | | | | | | | 0.0061 | 1 | | |
| DPA1*02:01:01 | 0.1383 | 283 | 0.1333 | 100 | 0.1154 | 78 | 0.1292 | 23 | 0.1646 | 27 | 0.1978 | 55 |
| DPA1*02:01:02 | 0.0024 | 5 | | | 0.0015 | 1 | | | 0.0061 | 1 | 0.0108 | 3 |
| DPA1*02:01:07 | 0.0098 | 20 | 0.0080 | 6 | 0.0059 | 4 | 0.0169 | 3 | 0.0183 | 3 | 0.0144 | 4 |
| DPA1*02:01:08 | 0.0670 | 137 | 0.0373 | 28 | 0.0828 | 56 | 0.0674 | 12 | 0.0854 | 14 | 0.0971 | 27 |
| DPA1*02:02:02 | 0.1711 | 350 | 0.1680 | 126 | 0.1598 | 108 | 0.1854 | 33 | 0.2317 | 38 | 0.1619 | 45 |
| DPA1*02:09:01 | 0.0103 | 21 | 0.0240 | 18 | 0.0044 | 3 | | | | | | |
| DPA1*02:09:02 | 0.0005 | 1 | | | | | | | 0.0061 | 1 | | |
| DPA1*02:12:01 | 0.0015 | 3 | | | | | | | | | 0.0108 | 3 |
| DPA1*02:34:01 | 0.0005 | 1 | 0.0013 | 1 | | | | | | | | |
| DPA1*02:89 | 0.0005 | 1 | 0.0013 | 1 | | | | | | | | |
| DPA1*03:01:01 | 0.1696 | 347 | 0.1547 | 116 | 0.2204 | 149 | 0.1292 | 23 | 0.1890 | 31 | 0.1007 | 28 |
| DPA1*03:01:02 | 0.0088 | 18 | 0.0080 | 6 | 0.0074 | 5 | 0.0056 | 1 | 0.0061 | 1 | 0.0180 | 5 |
| DPA1*03:01:03 | 0.0039 | 8 | 0.0040 | 3 | 0.0074 | 5 | | | | | | |
| DPA1*03:02 | 0.0010 | 2 | | | | | 0.0056 | 1 | | | 0.0036 | 1 |
| DPA1*03:05:01 | 0.0034 | 7 | 0.0013 | 1 | 0.0015 | 1 | | | 0.0061 | 1 | 0.0144 | 4 |
| DPA1*03:05:02Q | 0.0005 | 1 | | | 0.0015 | 1 | | | | | | |
| DPA1*04:02:01 | 0.0274 | 56 | 0.0280 | 21 | 0.0399 | 27 | 0.0281 | 5 | 0.0183 | 3 | | |
| DPA1*00:00:00 | 0.0025 | 5 | 0.0026 | 2 | 0.0030 | 2 | | | | | 0.0036 | 1 |

Overall allele frequencies at 3-field resolution for HLA-DPA1 locus were calculated per population.

The 2N values reflect total number of alleles found per HLA-DPA1 allele. "DPA1*00:00:00" reflects missing data which includes novel alleles awaiting further or final confirmation.

**Supplementary Table 2.** HLA-DQA1 allele frequencies in Africa across East, Southeast, and West countries at 3-field resolution

| | Africa | | East | | | | | | Southeast | | West | |
|---|---|---|---|---|---|---|---|---|---|---|---|---|
| | Overall | | Kenya | | Uganda | | Tanzania | | Mozambique | | Nigeria | |
| **Allele** | Frequency | 2N=2046 | Frequency | 2N=750 | Frequency | 2N=676 | Frequency | 2N=178 | Frequency | 2N=164 | Frequency | 2N=278 |
| DQA1*01:01:01 | 0.0132 | 27 | 0.0107 | 8 | 0.0178 | 12 | 0.0225 | 4 | 0.0122 | 2 | 0.0036 | 1 |
| DQA1*01:01:02 | 0.0763 | 156 | 0.0853 | 64 | 0.0917 | 62 | 0.0730 | 13 | 0.0305 | 5 | 0.0432 | 12 |
| DQA1*01:02:01 | 0.3201 | 655 | 0.2880 | 216 | 0.3373 | 228 | 0.3258 | 58 | 0.3293 | 54 | 0.3561 | 99 |
| DQA1*01:02:02 | 0.0039 | 8 | | | 0.0030 | 2 | 0.0056 | 1 | | | 0.0180 | 5 |
| DQA1*01:02:03 | 0.0005 | 1 | | | | | | | | | 0.0036 | 1 |
| DQA1*01:03:01 | 0.0425 | 87 | 0.0373 | 28 | 0.0281 | 19 | 0.0506 | 9 | 0.0915 | 15 | 0.0576 | 16 |
| DQA1*01:04:01 | 0.0049 | 10 | 0.0040 | 3 | 0.0059 | 4 | | | 0.0122 | 2 | 0.0036 | 1 |
| DQA1*01:05:01 | 0.0709 | 145 | 0.0547 | 41 | 0.0828 | 56 | 0.0955 | 17 | 0.0610 | 10 | 0.0755 | 21 |
| DQA1*01:05:02 | 0.0034 | 7 | 0.0053 | 4 | | | | | 0.0061 | 1 | 0.0072 | 2 |
| DQA1*01:05:04 | 0.0005 | 1 | | | 0.0015 | 1 | | | | | | |
| DQA1*01:36 | 0.0034 | 7 | 0.0027 | 2 | 0.0059 | 4 | 0.0056 | 1 | | | | |
| DQA1*01:63:01 | 0.0078 | 16 | 0.0173 | 13 | 0.0044 | 3 | | | | | | |
| DQA1*02:01:01 | 0.0557 | 114 | 0.0480 | 36 | 0.0695 | 47 | 0.0449 | 8 | 0.0366 | 6 | 0.0612 | 17 |
| DQA1*03:03:01 | 0.0670 | 137 | 0.0627 | 47 | 0.0562 | 38 | 0.0787 | 14 | 0.0671 | 11 | 0.0971 | 27 |
| DQA1*04:01:01 | 0.0738 | 151 | 0.0720 | 54 | 0.0799 | 54 | 0.0730 | 13 | 0.0915 | 15 | 0.0540 | 15 |
| DQA1*04:01:02 | 0.0235 | 48 | 0.0200 | 15 | 0.0237 | 16 | 0.0169 | 3 | 0.0183 | 3 | 0.0396 | 11 |
| DQA1*04:03N | 0.0005 | 1 | | | 0.0015 | 1 | | | | | | |
| DQA1*05:01:01 | 0.0665 | 136 | 0.0947 | 71 | 0.0399 | 27 | 0.0449 | 8 | 0.0671 | 11 | 0.0684 | 19 |
| DQA1*05:02 | 0.0039 | 8 | 0.0013 | 1 | 0.0044 | 3 | | | 0.0244 | 4 | | |
| DQA1*05:05:01 | 0.1579 | 323 | 0.1893 | 142 | 0.1435 | 97 | 0.1629 | 29 | 0.1524 | 25 | 0.1079 | 30 |
| DQA1*05:11:01 | 0.0015 | 3 | 0.0013 | 1 | 0.0030 | 2 | | | | | | |
| DQA1*00:00:00 | 0.0025 | 5 | 0.0054 | 2 | | | | | | | 0.0036 | 1 |

Overall allele frequencies at 3-field resolution for HLA-DQA1 locus were calculated per population.

The 2N values reflect total number of alleles found per HLA-DQA1 allele. "DQA1*00:00:00" reflects missing data which includes novel alleles awaiting official naming.

**Supplementary Table 3.** HLA-DRB3/4/5 allele frequencies in Africa across East, Southeast, and West countries at 3-field resolution

| | **Africa** | | **East** | | | | | | **Southeast** | | **West** | |
|---|---|---|---|---|---|---|---|---|---|---|---|---|
| | Overall | | Kenya | | Uganda | | Tanzania | | Mozambique | | Nigeria | |
| **DRB3** | Frequency | 2N=2046 | Frequency | 2N=750 | Frequency | 2N=676 | Frequency | 2N=178 | Frequency | 2N=164 | Frequency | 2N=278 |
| DRB3*00:00:00 | 0.4135 | 846 | 0.3947 | 296 | 0.4083 | 276 | 0.4607 | 82 | 0.3171 | 52 | 0.5036 | 140 |
| DRB3*01:01:02 | 0.0528 | 108 | 0.0554 | 41 | 0.0385 | 26 | 0.0562 | 10 | 0.0610 | 10 | 0.0755 | 21 |
| DRB3*01:62:01 | 0.0674 | 138 | 0.0613 | 46 | 0.0710 | 48 | 0.0730 | 13 | 0.0915 | 15 | 0.0576 | 16 |
| DRB3*02:02:01 | 0.3153 | 645 | 0.3187 | 239 | 0.3136 | 212 | 0.3090 | 55 | 0.3476 | 57 | 0.2950 | 82 |
| DRB3*02:02:03 | 0.0010 | 2 | 0.0027 | 2 | | | | | | | | |
| DRB3*02:71 | 0.0005 | 1 | 0.0013 | 1 | | | | | | | | |
| DRB3*03:01:01 | 0.1427 | 292 | 0.1547 | 116 | 0.1642 | 111 | 0.0899 | 16 | 0.1829 | 30 | 0.0683 | 19 |
| **DRB3*00:00:00** | 0.0069 | 14 | 0.0120 | 9 | 0.0044 | 3 | 0.0112 | 2 | | | | |

| | Overall | | Kenya | | Uganda | | Tanzania | | Mozambique | | Nigeria | |
|---|---|---|---|---|---|---|---|---|---|---|---|---|
| **DRB4** | Frequency | 2N=2046 | Frequency | 2N=750 | Frequency | 2N=676 | Frequency | 2N=178 | Frequency | 2N=164 | Frequency | 2N=278 |
| DRB4*00:00:00 | 0.8983 | 1838 | 0.8987 | 674 | 0.8994 | 608 | 0.8876 | 158 | 0.9146 | 150 | 0.8921 | 248 |
| DRB4*01:01:01 | 0.0362 | 74 | 0.0200 | 15 | 0.0355 | 24 | 0.0506 | 9 | 0.0488 | 8 | 0.0647 | 18 |
| DRB4*01:03:01 | 0.0645 | 132 | 0.0813 | 61 | 0.0636 | 43 | 0.0618 | 11 | 0.0366 | 6 | 0.0396 | 11 |
| DRB4*01:136 | 0.0005 | 1 | | | | | | | | | 0.0036 | 1 |
| DRB4*01:15 | 0.0005 | 1 | | | 0.0015 | 1 | | | | | | |

| | Overall | | Kenya | | Uganda | | Tanzania | | Mozambique | | Nigeria | |
|---|---|---|---|---|---|---|---|---|---|---|---|---|
| **DRB5** | Frequency | 2N=2046 | Frequency | 2N=750 | Frequency | 2N=676 | Frequency | 2N=178 | Frequency | 2N=164 | Frequency | 2N=278 |
| DRB5*00:00:00 | 0.8460 | 1731 | 0.8720 | 654 | 0.8476 | 573 | 0.8202 | 146 | 0.8720 | 143 | 0.7734 | 215 |
| DRB5*01:01:01 | 0.1481 | 303 | 0.1280 | 96 | 0.1494 | 101 | 0.1685 | 30 | 0.1159 | 19 | 0.2050 | 57 |
| DRB5*01:02:01 | 0.0005 | 1 | | | | | 0.0056 | 1 | | | | |
| DRB5*02:21:01 | 0.0024 | 5 | | | 0.0030 | 2 | 0.0056 | 1 | | | 0.0072 | 2 |
| DRB5*02:22 | 0.0005 | 1 | | | | | | | | | 0.0036 | 1 |
| **DRB5*00:00:00** | 0.0024 | 5 | | | | | | | 0.0122 | 2 | 0.0108 | 3 |

Overall allele frequencies at 3-field resolution for HLA-DRB3/4/5 were calculated per population. The frequency of each HLA-DRB3/4/5 allele is represented per country.
The 2N values reflect total number of alleles found per HLA-DRB3/4/5 allele. DRB3*00:00:00, DRB4*00:00:00, or DRB5*00:00:00 denotes unexpected based on DRB1 linkage association.
"**DRB3*00:00:00**" and "**DRB5*00:00:00**" reflects missing data which includes novel alleles awaiting official naming.

**Supplementary Table 4.** Various population genetics measurements in the African population

| Africa (n=1023) | | | | | | | | | | |
|---|---|---|---|---|---|---|---|---|---|---|
| | | Heterozygosity | | | Homozygosity | | | | Diversity | |
| Locus | Allele # | Obs.Het. | Exp.Het. | P-value | Obs.F. | Exp. F. | Fnd | P-value | Shannon | Simpson |
| A | 49 | 0.930 | 0.944 | 0.395 | 0.056 | 0.100 | -1.321 | 0.011* | 3.172 | 0.944 |
| B | 72 | 0.948 | 0.953 | 0.332 | 0.048 | 0.065 | -0.919 | 0.126 | 3.382 | 0.953 |
| C | 46 | 0.923 | 0.919 | 0.152 | 0.082 | 0.107 | -0.693 | 0.241 | 2.878 | 0.918 |
| DPA1 | 30 | 0.785 | 0.786 | 0.543 | 0.215 | 0.167 | 0.758 | 0.826 | 1.863 | 0.785 |
| DPB1 | 65 | 0.868 | 0.879 | 0.190 | 0.122 | 0.072 | 2.294 | 0.967 | 2.641 | 0.879 |
| DQA1 | 24 | 0.836 | 0.842 | 0.769 | 0.158 | 0.207 | -0.598 | 0.304 | 2.218 | 0.842 |
| DQB1 | 28 | 0.808 | 0.877 | 0.001** | 0.126 | 0.178 | -0.764 | 0.202 | 2.423 | 0.877 |
| DRB1 | 42 | 0.900 | 0.919 | 0.090 | 0.081 | 0.118 | -0.888 | 0.133 | 2.763 | 0.919 |
| DRB3 | 7 | 0.705 | 0.702 | 0.000** | 0.301 | 0.491 | -1.066 | 0.126 | | |
| DRB4 | 4 | 0.187 | 0.188 | 0.177 | 0.813 | 0.636 | 0.881 | 0.750 | | |
| DRB5 | 4 | 0.261 | 0.262 | 0.139 | 0.742 | 0.636 | 0.528 | 0.665 | | |

| Kenya (n=375) | | | | | | | | | | |
|---|---|---|---|---|---|---|---|---|---|---|
| | | Heterozygosity | | | Homozygosity | | | | Diversity | |
| Locus | Allele # | Obs.Het. | Exp.Het. | P-value | Obs.F. | Exp. F. | Fnd | P-value | Shannon | Simpson |
| A | 43 | 0.925 | 0.939 | 0.455 | 0.063 | 0.092 | -1.044 | 0.077 | 3.150 | 0.938 |
| B | 56 | 0.949 | 0.961 | 0.187 | 0.041 | 0.067 | -1.424 | 0.007* | 3.465 | 0.959 |
| C | 39 | 0.952 | 0.923 | 0.315 | 0.078 | 0.104 | -0.762 | 0.201 | 2.923 | 0.922 |
| DPA1 | 20 | 0.760 | 0.770 | 0.463 | 0.231 | 0.209 | 0.272 | 0.708 | 1.824 | 0.769 |
| DPB1 | 39 | 0.867 | 0.891 | 0.003* | 0.111 | 0.104 | 0.208 | 0.690 | 2.641 | 0.890 |
| DQA1 | 19 | 0.856 | 0.849 | 0.964 | 0.152 | 0.220 | -0.805 | 0.186 | 2.209 | 0.848 |
| DQB1 | 21 | 0.765 | 0.887 | 0.000** | 0.117 | 0.199 | -1.075 | 0.059 | 2.441 | 0.886 |
| DRB1 | 23 | 0.904 | 0.917 | 0.105 | 0.084 | 0.182 | -1.426 | 0.003* | 2.669 | 0.916 |
| DRB3 | 6 | 0.741 | 0.713 | 0.001** | 0.294 | 0.492 | -1.119 | 0.107 | 3.150 | 0.938 |
| DRB4 | 2 | 0.192 | 0.186 | 0.771 | 0.815 | 0.754 | 0.313 | 0.528 | | |
| DRB5 | 1 | 0.219 | 0.224 | 0.644 | 0.777 | 0.862 | -0.517 | 0.268 | | |

| Uganda (n=338) | | | | | | | | | | |
|---|---|---|---|---|---|---|---|---|---|---|
| Locus | Allele # | Obs.Het. | Exp.Het. | P-value | Obs.F. | Exp. F. | Fnd | P-value | Shannon | Simpson |
| A | 37 | 0.941 | 0.942 | 0.427 | 0.059 | 0.107 | -1.368 | 0.006* | 3.085 | 0.941 |
| B | 51 | 0.950 | 0.945 | 0.379 | 0.056 | 0.073 | -0.815 | 0.172 | 3.185 | 0.944 |
| C | 34 | 0.923 | 0.915 | 0.774 | 0.086 | 0.118 | -0.803 | 0.175 | 2.795 | 0.914 |
| DPA1 | 18 | 0.817 | 0.788 | 0.534 | 0.213 | 0.227 | -0.162 | 0.543 | 1.797 | 0.787 |
| DPB1 | 39 | 0.882 | 0.863 | 0.262 | 0.139 | 0.101 | 1.170 | 0.895 | 2.496 | 0.862 |
| DQA1 | 19 | 0.796 | 0.834 | 0.333 | 0.167 | 0.216 | -0.589 | 0.309 | 2.171 | 0.833 |
| DQB1 | 21 | 0.846 | 0.867 | 0.094 | 0.135 | 0.196 | -0.821 | 0.171 | 2.318 | 0.866 |
| DRB1 | 29 | 0.902 | 0.915 | 0.590 | 0.086 | 0.140 | -1.094 | 0.054 | 2.708 | 0.914 |
| DRB3 | 4 | 0.701 | 0.702 | 0.215 | 0.300 | 0.593 | -1.531 | 0.028 | | |
| DRB4 | 3 | 0.186 | 0.186 | 0.865 | 0.814 | 0.665 | 0.757 | 0.703 | | |
| DRB5 | 2 | 0.257 | 0.260 | 0.875 | 0.741 | 0.751 | -0.053 | 0.449 | | |

| Tanzania (n=89) | | | | | | | | | | |
|---|---|---|---|---|---|---|---|---|---|---|
| Locus | Allele # | Obs.Het. | Exp.Het. | P-value | Obs.F. | Exp. F. | Fnd | P-value | Shannon | Simpson |
| A | 29 | 0.933 | 0.936 | 0.522 | 0.069 | 0.097 | -0.981 | 0.094 | 2.919 | 0.931 |
| B | 32 | 0.966 | 0.933 | 0.134 | 0.073 | 0.085 | -0.537 | 0.323 | 2.931 | 0.927 |
| C | 19 | 0.910 | 0.891 | 0.973 | 0.114 | 0.161 | -0.845 | 0.154 | 2.437 | 0.886 |
| DPA1 | 11 | 0.742 | 0.753 | 0.883 | 0.251 | 0.282 | -0.300 | 0.469 | 1.673 | 0.749 |
| DPB1 | 22 | 0.899 | 0.877 | 0.247 | 0.128 | 0.135 | -0.152 | 0.537 | 2.409 | 0.872 |
| DQA1 | 13 | 0.820 | 0.839 | 0.149 | 0.166 | 0.239 | -0.854 | 0.170 | 2.110 | 0.834 |
| DQB1 | 17 | 0.809 | 0.856 | 0.162 | 0.149 | 0.181 | -0.509 | 0.354 | 2.226 | 0.851 |
| DRB1 | 22 | 0.865 | 0.918 | 0.085 | 0.087 | 0.135 | -1.087 | 0.056 | 2.705 | 0.913 |
| DRB3 | 5 | 0.640 | 0.679 | 0.037* | 0.324 | 0.465 | -0.886 | 0.201 | | |
| DRB4 | 2 | 0.157 | 0.207 | 0.006* | 0.794 | 0.701 | 0.498 | 0.616 | | |
| DRB5 | 3 | 0.303 | 0.300 | 0.403 | 0.701 | 0.604 | 0.529 | 0.677 | | |

| Mozambique (n=82) | | | | | | | | | | |
|---|---|---|---|---|---|---|---|---|---|---|
| Locus | Allele # | Obs.Het. | Exp.Het. | P-value | Obs.F. | Exp. F. | Fnd | P-value | Shannon | Simpson |
| A | 25 | 0.939 | 0.941 | 0.131 | 0.065 | 0.113 | -1.429 | 0.005* | 2.922 | 0.936 |
| B | 29 | 0.951 | 0.946 | 0.725 | 0.060 | 0.094 | -1.274 | 0.017* | 3.009 | 0.940 |
| C | 23 | 0.915 | 0.924 | 0.662 | 0.081 | 0.125 | -1.117 | 0.049 | 2.763 | 0.919 |
| DPA1 | 13 | 0.805 | 0.818 | 0.862 | 0.187 | 0.235 | -0.570 | 0.320 | 1.863 | 0.813 |
| DPB1 | 25 | 0.817 | 0.847 | 0.142 | 0.158 | 0.113 | 1.322 | 0.905 | 2.340 | 0.842 |
| DQA1 | 14 | 0.854 | 0.840 | 0.856 | 0.165 | 0.218 | -0.684 | 0.255 | 2.153 | 0.835 |
| DQB1 | 14 | 0.866 | 0.871 | 0.610 | 0.134 | 0.218 | -1.080 | 0.061 | 2.238 | 0.866 |
| DRB1 | 22 | 0.890 | 0.910 | 0.784 | 0.095 | 0.132 | -0.866 | 0.146 | 2.586 | 0.905 |
| DRB3 | 4 | 0.768 | 0.738 | 0.220 | 0.267 | 0.522 | -1.492 | 0.020* | | |
| DRB4 | 2 | 0.171 | 0.161 | 1.000 | 0.840 | 0.697 | 0.766 | 0.692 | | |
| DRB5 | 1 | 0.232 | 0.228 | 1.000 | 0.791 | 0.825 | -0.204 | 0.356 | | |

| Nigeria (n=139) | | | | | | | | | | |
|---|---|---|---|---|---|---|---|---|---|---|
| Locus | Allele # | Obs.Het. | Exp.Het. | P-value | Obs.F. | Exp. F. | Fnd | P-value | Shannon | Simpson |
| A | 30 | 0.906 | 0.941 | 0.171 | 0.062 | 0.107 | -1.339 | 0.010* | 2.991 | 0.938 |
| B | 36 | 0.928 | 0.936 | 0.880 | 0.068 | 0.086 | -0.733 | 0.215 | 3.047 | 0.932 |
| C | 29 | 0.856 | 0.903 | 0.046 | 0.100 | 0.112 | -0.328 | 0.450 | 2.725 | 0.900 |
| DPA1 | 14 | 0.791 | 0.793 | 0.822 | 0.210 | 0.245 | -0.375 | 0.435 | 1.844 | 0.790 |
| DPB1 | 26 | 0.849 | 0.870 | 0.372 | 0.133 | 0.127 | 0.154 | 0.672 | 2.464 | 0.867 |
| DQA1 | 16 | 0.878 | 0.831 | 0.718 | 0.172 | 0.215 | -0.542 | 0.333 | 2.158 | 0.828 |
| DQB1 | 18 | 0.799 | 0.859 | 0.066 | 0.150 | 0.189 | -0.572 | 0.317 | 2.366 | 0.856 |
| DRB1 | 27 | 0.914 | 0.910 | 0.738 | 0.093 | 0.122 | -0.730 | 0.217 | 2.713 | 0.907 |
| DRB3 | 4 | 0.619 | 0.648 | 0.455 | 0.353 | 0.551 | -1.100 | 0.129 | | |
| DRB4 | 3 | 0.201 | 0.199 | 0.112 | 0.802 | 0.627 | 0.923 | 0.760 | | |
| DRB5 | 3 | 0.374 | 0.361 | 0.436 | 0.656 | 0.626 | 0.159 | 0.581 | | |

**Heterozygosity** index was measured using Hardy-Weinberg Equilibrium (HWE) for each population in Arlequin for all loci using Markov chain. The p-value for HWE deviation (p-HWE) reported as *p<0.05 & **p<0.001 reflects the difference between observed heterozygosity (Obs. Het.) and expected heterozygosity (Exp. Het.). **Homozygosity** index was measured using Slatkin's implementation of Ewens-Watterson (EW) homozygosity test of neutrality. Observed homozygosity F (Obs. F) is the sum of the squared allele frequencies from the mean value of Expected homozygosity F (Exp. F) for a population (2n) with the same number of unique alleles undergoing neutral evolution. For each locus the normalized deviate of F (Fnd) is the difference between the Obs. F and Exp. F divided by the square root of the variance of the expected homozygosity. Significant negative Fnd values (<0.025) correspond to lower observed F values than expected mean F values. The p-value for EW reported as *p<0.05 & **p<0.001 . **Diversity** index was measured using Shannon (richness and evenness of alleles relative to frequency) and Simpson (weighted abundance of alleles relative to frequency).

**Supplementary Table 5.** Multi-locus haplotypes in the overall African population with frequencies <5% and >5% within individual populations.

| | Africa | East | | | Southeast | West |
|---|---|---|---|---|---|---|
| | Overall | Kenya | Uganda | Tanzania | Mozambique | Nigeria |
| Haplotype | 2N=2046 | 2N=750 | 2N=676 | 2N=178 | 2N=164 | 2N=278 |
| DPB1*02:01:02~DQB1*06:02:01 | 0.0497 | 0.0243 | 0.0642 | 0.0726 | 0.0305 | 0.0931 |
| DQA1*05:05:01~DQB1*03:19:01 | 0.0494 | 0.0474 | 0.0504 | 0.0787 | 0.1098 | 0.0074 |
| DQA1*02:01:01~DQB1*02:02:01 | 0.0479 | 0.0393 | 0.0593 | 0.0393 | 0.0366 | 0.0551 |
| DQB1*02:02:01~DRB1*07:01:01 | 0.0464 | 0.0420 | 0.0460 | 0.0449 | 0.0549 | 0.0549 |
| DQA1*01:02:01~DQB1*06:09:01 | 0.0464 | 0.0618 | 0.0534 | 0.0169 | 0.0406 | 0.0112 |
| DQB1*06:09:01~DRB1*13:02:01 | 0.0462 | 0.0623 | 0.0534 | 0.0169 | 0.0427 | 0.0074 |
| DPA1*02:01:01~DPB1*17:01:01 | 0.0455 | 0.0481 | 0.0475 | 0.0225 | 0.0122 | 0.0683 |
| DPB1*01:01:01~DQB1*05:01:01 | 0.0454 | 0.0278 | 0.0577 | 0.0636 | 0.0521 | 0.0703 |
| DPB1*01:01:01~DRB1*11:01:02 | 0.0443 | 0.0460 | 0.0417 | 0.0787 | 0.0629 | 0.0186 |
| DRB1*08:04:01~DRB3/4/5*00:00:00 | 0.0441 | 0.0447 | 0.0357 | 0.0225 | 0.0188 | 0.0919 |
| DPA1*02:02:02~DPB1*01:01:01~DQA1*04:01:01~DQB1*04:02:01~DRB1*03:02:01~DRB3*01:62:01 | 0.0439 | 0.0364 | 0.0490 | 0.0441 | 0.0813 | 0.0292 |
| DPA1*01:03:01~DPB1*104:01:01 | 0.0431 | 0.0722 | 0.0148 | 0.0506 | 0.0427 | 0.0288 |
| C*04:01:01~DRB1*11:01:02 | 0.0423 | 0.0259 | 0.0590 | 0.0832 | 0.0145 | 0.0495 |
| DQA1*01:02:01~DQB1*06:04:01 | 0.0421 | 0.0537 | 0.0492 | 0.0281 | 0.0305 | 0.0074 |
| DQB1*06:04:01~DRB1*13:02:01 | 0.0416 | 0.0528 | 0.0504 | 0.0281 | 0.0305 | 0.0037 |
| DRB1*11:02:01~DRB3*02:02:01 | 0.0416 | 0.0362 | 0.0585 | 0.0359 | 0.0316 | 0.0257 |
| DPB1*105:01:01~DRB1*11:01:02 | 0.0410 | 0.0367 | 0.0641 | 0.0122 | 0.0393 | 0.0167 |
| B*15:10:01~C*03:04:02 | 0.0406 | 0.0254 | 0.0429 | 0.0955 | 0.0427 | 0.0396 |
| DQB1*03:01:01~DRB1*11:02:01 | 0.0398 | 0.0406 | 0.0606 | 0.0276 | 0.0122 | 0.0074 |
| DRB1*10:01:01~DRB3/4/5*00:00:00 | 0.0396 | 0.0420 | 0.0372 | 0.0506 | 0.0438 | 0.0294 |
| DPA1*01:03:01~DPB1*02:01:02~DQA1*01:02:01~DQB1*06:02:01~DRB1*15:03:01~DRB5*01:01:01 | 0.0393 | 0.0210 | 0.0541 | 0.0601 | 0.0250 | 0.0677 |
| DPA1*03:01:01~DPB1*105:01:01~DQA1*01:02:01~DQB1*06:02:01~DRB1*15:03:01~DRB5*01:01:01 | 0.0392 | 0.0332 | 0.0354 | 0.0377 | 0.0687 | 0.0451 |
| DQB1*05:01:01~DRB1*10:01:01 | 0.0390 | 0.0420 | 0.0371 | 0.0506 | 0.0427 | 0.0257 |
| DPB1*02:01:02~DRB1*15:03:01 | 0.0386 | 0.0216 | 0.0527 | 0.0571 | 0.0291 | 0.0592 |
| DPB1*105:01:01~DRB1*15:03:01 | 0.0366 | 0.0300 | 0.0327 | 0.0379 | 0.0668 | 0.0342 |
| B*45:01:01~C*16:01:01 | 0.0362 | 0.0307 | 0.0444 | 0.0674 | 0.0183 | 0.0216 |
| DQB1*03:19:01~DRB1*11:01:02 | 0.0358 | 0.0402 | 0.0328 | 0.0500 | 0.0671 | |
| B*07:02:01~C*07:02:01 | 0.0356 | 0.0281 | 0.0355 | 0.0674 | 0.0061 | 0.0540 |
| DPB1*01:01:01~DQB1*06:02:01 | 0.0350 | 0.0207 | 0.0294 | 0.0494 | 0.0448 | 0.0663 |
| B*58:01:01~C*07:18:01 | 0.0337 | 0.0320 | 0.0191 | 0.0281 | 0.0671 | 0.0576 |
| DRB1*13:01:01~DRB3*02:02:01 | 0.0332 | 0.0341 | 0.0226 | 0.0300 | 0.0438 | 0.0535 |
| C*04:01:01~DRB1*15:03:01 | 0.0318 | 0.0277 | 0.0290 | 0.0064 | 0.0388 | 0.0556 |
| DPB1*01:01:01~DQB1*03:19:01 | 0.0289 | 0.0350 | 0.0234 | 0.0513 | 0.0493 | |
| A*36:01:01~C*04:01:01 | 0.0278 | 0.0120 | 0.0233 | 0.0895 | 0.0183 | 0.0468 |
| DQA1*03:03:01~DQB1*02:02:01 | 0.0276 | 0.0122 | 0.0282 | 0.0449 | 0.0366 | 0.0515 |
| A*02:01:01~C*16:01:01 | 0.0272 | 0.0260 | 0.0283 | 0.0506 | | 0.0386 |
| DPA1*02:02:02~DPB1*01:01:01~DQA1*05:05:01~DQB1*03:19:01~DRB1*11:01:02~DRB3*02:02:01 | 0.0268 | 0.0341 | 0.0218 | 0.0506 | 0.0375 | |
| A*36:01:01~B*53:01:01~C*04:01:01 | 0.0261 | 0.0121 | 0.0206 | 0.0838 | 0.0244 | 0.0432 |
| A*36:01:01~B*53:01:01 | 0.0259 | 0.0121 | 0.0183 | 0.0831 | 0.0244 | 0.0428 |
| DPB1*18:01:01~DQB1*05:01:01 | 0.0255 | 0.0136 | 0.0403 | 0.0746 | 0.0122 | 0.0124 |
| DPB1*105:01:01~DQB1*05:01:01 | 0.0245 | 0.0133 | 0.0528 | 0.0215 | 0.0067 | 0.0082 |
| B*53:01:01~DRB1*11:01:02 | 0.0241 | 0.0175 | 0.0190 | 0.0660 | 0.0071 | 0.0310 |
| C*06:02:01~DRB1*15:03:01 | 0.0218 | 0.0228 | 0.0231 | 0.0540 | 0.0260 | |
| DPA1*02:01:01~DPB1*13:01:01 | 0.0215 | 0.0107 | 0.0148 | 0.0393 | 0.0854 | 0.0180 |
| A*36:01:01~DRB1*11:01:02 | 0.0200 | 0.0099 | 0.0164 | 0.0787 | 0.0061 | 0.0213 |
| DPB1*01:01:01~DRB1*15:03:01 | 0.0188 | 0.0212 | 0.0125 | 0.0144 | 0.0139 | 0.0531 |
| DQA1*01:03:01~DQB1*06:03:01 | 0.0178 | 0.0108 | 0.0119 | 0.0225 | 0.0671 | 0.0184 |
| DQB1*06:03:01~DRB1*13:01:01 | 0.0177 | 0.0107 | 0.0119 | 0.0225 | 0.0671 | 0.0184 |
| DRB1*11:02:01~DRB3*03:01:01 | 0.0173 | 0.0153 | 0.0159 | 0.0146 | 0.0622 | |
| A*68:02:01~C*03:04:02 | 0.0153 | 0.0126 | 0.0168 | 0.0501 | | 0.0084 |
| DQB1*03:19:01~DRB1*11:02:01 | 0.0153 | 0.0080 | 0.0136 | 0.0230 | 0.0671 | 0.0037 |
| A*02:01:01~DRB1*15:03:01 | 0.0143 | 0.0292 | | 0.0506 | 0.0106 | 0.0094 |
| A*68:02:01~B*15:10:01~C*03:04:02 | 0.0140 | 0.0083 | 0.0130 | 0.0501 | | 0.0081 |
| A*02:05:01~B*58:01:01 | 0.0112 | 0.0106 | | 0.0225 | 0.0549 | 0.0072 |
| A*02:05:01~C*07:18:01 | 0.0104 | 0.0062 | 0.0015 | 0.0169 | 0.0610 | 0.0072 |
| B*44:03:02~C*07:06:01 | 0.0093 | 0.0027 | 0.0015 | | 0.0732 | 0.0144 |
| A*02:05:01~B*58:01:01~C*07:18:01 | 0.0092 | 0.0065 | | 0.0169 | 0.0549 | 0.0072 |
| DPB1*13:01:01~DRB1*11:02:01 | 0.0068 | | 0.0033 | 0.0169 | 0.0598 | |

Multi-locus haplotypes in the African population with frequencies <5% in the overall and >5% within individual populations were calculated using Hapl-o-mat v1.3.

**Supplementary Table 6.** Unique HLA allele frequencies in Africa across East, Southeast, and West countries at 3-field resolution

| East | | | | | | Southeast | | West | |
|---|---|---|---|---|---|---|---|---|---|
| Kenya | | Uganda | | Tanzania | | Mozambique | | Nigeria | |
| **Allele** | 2N=750 | **Allele** | 2N=676 | **Allele** | 2N=178 | **Allele** | 2N=164 | **Allele** | 2N=278 |
| A*02:02PN | 1 | A*01:23:01 | 1 | B*38:01:01 | 1 | A*29:11 | 1 | A*02:60:01 | 2 |
| A*02:85 | 1 | A*02:951 | 1 | DQB1*06:01:01 | 1 | B*15:01:01 | 1 | A*30:02PN | 1 |
| A*26:30 | 4 | B*15:610PN | 1 | DRB1*15:02:01 | 1 | C*03:03:01 | 2 | A*74:09 | 1 |
| A*29:15 | 4 | B*27:05:02 | 1 | DRB1*15:03PN | 1 | DPA1*01:58:01 | 1 | B*14:03 | 3 |
| A*32:106:01 | 1 | B*35:03:01 | 1 | DRB3*02:02:01PN | 2 | DPA1*02:09:02 | 1 | B*40:02:01 | 1 |
| A*68:278 | 2 | B*53:08:01 | 1 | DRB5*01:02:01 | 1 | DPB1*02:01:19PN | 1 | B*44:07 | 2 |
| B*07:06:01 | 1 | C*06:22 | 1 | | | DPB1*1049:01 | 1 | B*52:01:02 | 6 |
| B*15:64:02 | 1 | C*07:01:113 | 1 | | | DPB1*1069:01 | 1 | B*53:37 | 1 |
| B*35:189 | 1 | DPA1*01:12:01 | 2 | | | DPB1*23:01:01 | 1 | B*57:05 | 1 |
| B*39:01:01 | 1 | DPA1*03:01PN2 | 1 | | | DPB1*417:01:01 | 1 | B*78:01:01 | 3 |
| B*39:06:02 | 1 | DPA1*03:05:02Q | 1 | | | DPB1*654:01 | 1 | B*82:01:01 | 1 |
| B*57:01:01 | 9 | DPB1*02:01:02PN2 | 1 | | | DPB1*762:01 | 1 | C*03:03:04 | 1 |
| B*58:01PN | 1 | DPB1*40:01PN | 1 | | | DQB1*04:04 | 1 | C*07:01PN | 1 |
| B*58:64 | 1 | DPB1*49:01PN | 1 | | | | | C*07:241 | 1 |
| C*04:290 | 1 | DPB1*61:01N | 3 | | | | | C*07:621 | 1 |
| C*07:18:11 | 1 | DPB1*71:01:01 | 1 | | | | | DPA1*01:19 | 2 |
| C*07:607 | 2 | DPB1*105:01:02 | 1 | | | | | DPA1*02:01:01PN | 1 |
| C*16:02:01 | 3 | DPB1*348:01:02 | 1 | | | | | DPA1*02:12:01 | 3 |
| C*16:04:01 | 2 | DPB1*1171:01 | 1 | | | | | DPB1*04:02:01 | 2 |
| DPA1*01:03:10 | 1 | DPB1*1325:01N | 1 | | | | | DPB1*10:01:01 | 1 |
| DPA1*01:06:02 | 1 | DQA1*01:05:04 | 1 | | | | | DPB1*1072:01 | 2 |
| DPA1*01:30 | 9 | DQA1*04:03N | 1 | | | | | DPB1*133:01:01 | 2 |
| DPA1*02:09PN | 1 | DQB1*02:70 | 1 | | | | | DPB1*584:01:01 | 2 |
| DPA1*02:34:01 | 1 | DQB1*04:52 | 1 | | | | | DPB1*63:01 | 1 |
| DPA1*02:89 | 1 | DQB1*04:87 | 3 | | | | | DPB1*85:01:01 | 3 |
| DPB1*02:01:22 | 9 | DRB1*01:01:01PN | 1 | | | | | DQA1*01:02:03 | 1 |
| DPB1*11:01:07 | 1 | DRB1*04:10:01 | 1 | | | | | DQA1*01:02PN2 | 1 |
| DPB1*19:01:01 | 1 | DRB1*08:08 | 3 | | | | | DQB1*05:186 | 1 |
| DPB1*88:01 | 2 | DRB1*11:01PN | 1 | | | | | DRB1*03:01:02 | 1 |
| DPB1*124:01:01 | 1 | DRB1*14:01:02 | 1 | | | | | DRB1*08:06:01 | 1 |
| DPB1*135:01PN | 1 | DRB4*01:15 | 1 | | | | | DRB1*11:04:02 | 2 |
| DPB1*233:01 | 2 | | | | | | | DRB1*11:08:02 | 1 |
| DPB1*463:01:01 | 1 | | | | | | | DRB1*11:10:01 | 2 |
| DPB1*495:01 | 1 | | | | | | | DRB1*13:01:03 | 1 |
| DPB1*1037:01 | 1 | | | | | | | DRB4*01:136 | 1 |
| DPB1*1428:01 | 1 | | | | | | | DRB5*02:22 | 1 |
| DQA1*01:02PN | 2 | | | | | | | | |
| DQA1*01:05:01PN | 2 | | | | | | | | |
| DQB1*06:02:23 | 1 | | | | | | | | |
| DQB1*06:14:02 | 1 | | | | | | | | |
| DQB1*06:85 | 1 | | | | | | | | |
| DRB1*03:01PN | 1 | | | | | | | | |
| DRB1*04:04:01 | 3 | | | | | | | | |
| DRB1*04:10PN | 1 | | | | | | | | |
| DRB3*02:02:03 | 2 | | | | | | | | |
| DRB3*02:71 | 1 | | | | | | | | |

Unique allele occurrence per population for all HLA alleles at 3-field resolution.

**Supplementary Table 7.** More recently identified novel alleles in the African cohorts

| Locus | allele | nucleotide change | amino acid change | exon | occurrence | Population | | New allele | IMGT/HLA accession number | Date Assigned |
|---|---|---|---|---|---|---|---|---|---|---|
| A | 02:05:01 | 790 A>G | 240 T>A | 4 | 1 | Uganda | | 02:951 | HLA29215 | 10/30/2020 |
| A | 68:02:01 | 806 T>C | 245 V>A | 4 | 2 | Kenya | | 68:278 | HLA32101 | 7/30/2021 |
| C | 07:01:05 | 912 C>T | 208 - | 5 | 1 | Uganda | | 07:01:113 | HLA35146 | 6/30/2022 |
| C | 07:18:01 | 123 C>A | 17 - | 2 | 1 | Kenya | | 07:18:11 | HLA37710 | 6/30/2023 |
| DPA1 | 02:01:11 | 319 C>T | 76 R>C | 2 | 1 | Mozambique | | 02:09:02 | HLA37155 | 2/28/2023 |
| DPA1 | 03:01:01 | 444 C>G | 117 - | 3 | 7 | Kenya | | 03:01:03 | HLA31052 | 5/28/2021 |
| DPA1 | 03:01:02 | 2 T>C | -31 M>T | 1 | 1 | Uganda | | 03:05:02Q | HLA34756 | 5/31/2022 |
| DPB1 | 105:01:01:01 | 739 C>T | 218 H>Y | 4 | 1 | Uganda | | 1171:01 | HLA30067 | 2/26/2021 |
| DQA1 | 01:01:01 | 169 G>C | 34 E>Q | 2 | 1 | Kenya | | 01:63:01 | HLA30382 | 2/26/2021 |
| DQA1 | 01:08 | 470 C>G | 134 A>G | 3 | 12 | Kenya | Uganda | 01:63:01 | HLA30382 | 2/26/2021 |
| DQA1 | 01:02:01 | 329 A>C | 87 N>T | 2 | 5 | Kenya | Uganda | 01:36 | HLA24078 | 6/28/2019 |
| DQA1 | 01:36 | * | - | - | 5 | Kenya | Uganda | - | HLA24078 | 6/28/2019 |
| DQB1 | 04:02:01 | 79 A>G | -6 T>A | 1 | 2 | Uganda | | 04:87 | HLA32159 | 7/30/2021 |
| DRB1 | 14:54:01 | 40 G>T | -16 V>F | 1 | 2 | Mozambique | Uganda | 14:243 | HLA34401 | 3/31/2022 |
| DRB4 | 01:01:01 | 577 G>A | 164 V>I | 3 | 1 | Nigeria | | 01:136 | HLA28875 | 9/30/2020 |

Novel alleles now validated by others in the IMGT/HLA database within the last five years. Listed are alleles which were placed in the novel allele category during the course of our NGS HLA typing of the African cohorts. * Allele is not novel, but IMGT currently contains only exon 2 and we are able to submit fully phased CDS and intronic sequence.

**Supplementary table 8.** Comparison with other published African population HLA studies*

| Region | Country | Study (Year) | Sample size | Resolution reported | Technology | Coverage | Genotyping software | IPD-IMGT HLA Database | class I alleles | | | class II alleles | | | | | | | |
|---|---|---|---|---|---|---|---|---|---|---|---|---|---|---|---|---|---|---|---|
| | | | | | | | | | HLA-A | HLA-B | HLA-C | DPA1 | DPB1 | DQA1 | DQB1 | DRB1 | DRB3 | DRB4 | DRB5 |
| East | Kenya | Our study | 375 | 2-field | NGS | Full | Omixon Target v1.8.0-1.9.3, Omixon HLA Explore v2.0 Explore, and NGSengine GenDX v2.10.0-3.30.0 | v.3.10-3.58 | 41 | 55 | 33 | 14 | 34 | 15 | 18 | 22 | 5 | 2 | 1 |
| East | Kenya | Banjoko et al., 2025 | 109 | 2-field | NGS | Exon | Omixon HLA Explore | v. 3.56 | 25 | 37 | 22 | - | - | - | - | - | - | - | - |
| East | Kenya | Arlehamn et al., 2017 | 100 | 2-field | NGS | Exon | In-house accredited HLA allele caller software pipeline | v.3.21.0 | 29 | 33 | 26 | - | 16 | 7 | 14 | 18 | 3 | 1 | 1 |
| East | Uganda | Our study | 338 | 2-field | NGS | Full | Omixon Target v1.8.0-1.9.3, Omixon HLA Explore v2.0 Explore, and NGSengine GenDX v2.10.0-3.30.0 | v.3.10-3.58 | 36 | 50 | 29 | 11 | 34 | 15 | 20 | 25 | 4 | 3 | 2 |
| East | Uganda | Digital et al., 2021 | 892 | 2-field | NGS | Exon | Omixon HLA Explore | - | 41 | 53 | 30 | 9 | 38 | 13 | 16 | 26 | - | - | - |
| East | Tanzania | Our study | 89 | 2-field | NGS | Full | Omixon Target v1.8.0-1.9.3, Omixon HLA Explore v2.0 Explore, and NGSengine GenDX v2.10.0-3.30.0 | v.3.10-3.58 | 27 | 32 | 18 | 7 | 19 | 10 | 16 | 22 | 4 | 2 | 3 |
| East | Tanzania | Barton et al., 2022 | 336 | 2-field | NGS | | Wheeler Aligner, xHLA using DIAMOND protein level aligner | - | 38 | 53 | 78 | - | 30 | - | 17 | 26 | - | - | - |
| Southeast | Mozambique | Our study | 82 | 2-field | NGS | Full | Omixon Target v1.8.0-1.9.3, Omixon HLA Explore v2.0 Explore, and NGSengine GenDX v2.10.0-3.30.0 | v.3.10-3.58 | 24 | 28 | 22 | 9 | 23 | 11 | 14 | 22 | 4 | 2 | 1 |
| West | Nigeria | Our study | 139 | 2-field | NGS | Full | Omixon Target v1.8.0-1.9.3, Omixon HLA Explore v2.0 Explore, and NGSengine GenDX v2.10.0-3.30.0 | v.3.10-3.58 | 30 | 35 | 26 | 9 | 24 | 11 | 16 | 24 | 4 | 3 | 3 |
| West | Nigeria | Testi et al., 2015 | 97 | 2-field | SSOP/SBT | Exon | - | - | 25 | 31 | 22 | - | - | - | 16 | 22 | - | - | - |

* studies include those typed with NGS and at least sequence-based 2-field data (n >50)
